# Parallel Hippocampal-Parietal Circuits for Self- and Goal-oriented Processing

**DOI:** 10.1101/2020.12.01.395210

**Authors:** Annie Zheng, David F. Montez, Scott Marek, Adrian W. Gilmore, Dillan J. Newbold, Timothy O. Laumann, Benjamin P. Kay, Nicole A. Seider, Andrew N. Van, Jacqueline M. Hampton, Dimitrios Alexopolous, Bradley L. Schlaggar, Chad M. Sylvester, Deanna J. Greene, Joshua S. Shimony, Steven M. Nelson, Gagan S. Wig, Caterina Gratton, Kathleen B. McDermott, Marcus E. Raichle, Evan M. Gordon, Nico U.F. Dosenbach

## Abstract

The hippocampus is critically important for a diverse range of cognitive processes, such as episodic memory, prospective memory, affective processing, and spatial navigation. The human hippocampus has been thought of as being solely functionally connected to a set of neocortical regions known as the default mode network (DMN), which supports self-referential cognition. Using individual-specific precision functional mapping of resting state fMRI data, we found the anterior hippocampus (head and body) to be preferentially connected to the DMN as expected. The hippocampal tail, however, was strongly preferentially connected to the parietal memory network (PMN), which supports goal-oriented cognition and stimulus recognition. This resting state functional connectivity (RSFC) anterior-posterior dichotomy was well-matched by differences in task deactivations and anatomical segmentations of the hippocampus. Task deactivations were localized to the head and body of the hippocampus (DMN), relatively sparing the tail (PMN). Anterior and posterior hippocampal connectivity was network-specific even though the DMN and PMN are interdigitated in medial parietal cortex. The functional dichotomization of the hippocampus into anterior DMN-connected and posterior PMN-connected parcels suggests parallel, but distinct circuits between the hippocampus and medial parietal cortex for self vs. goal-oriented processing.

## INTRODUCTION

The hippocampus is critically important for a diverse range of cognitive processes, such as episodic memory, prospective memory, affective processing, and spatial navigation (*1*–*8*). The hippocampus’ diverse functions rely on its pattern of connectivity, where episodic memories become transformed from being dependent on the hippocampus to being represented in the neocortex (*9*). Atypical cortico-hippocampal functional connectivity is associated with cognitive and affective deficits (*10*–*13*). A precise understanding of the functional organization of the hippocampus is crucial for understanding the neurobiology underlying hippocampally-related diseases.

The hippocampus seems to exhibit some functional heterogeneity. Studies of the rodent hippocampus have demonstrated modular differentiation along its longitudinal axis in patterns of gene expression, function, and anatomical projections (*2*, *14*, *15*). The rodent dorsal hippocampus (homologue of human posterior hippocampus) has been shown to be important for spatial navigation, whereas the ventral hippocampus (anterior hippocampus homologue) plays a role in the modulation of stress and affect (*2*, *4*). Hippocampal place field representation sizes in rodent models are also known to follow a dorsal-ventral gradient reflecting small-to-large spatial resolution (*14*, *15*). Anatomically, connections between entorhinal cortex and the hippocampus are arranged topographically along the dorsal-ventral axis (*4*). The ventral hippocampus in rats is interconnected with the amygdala, temporal pole, and ventromedial prefrontal cortex (*4*, *16*), while the dorsal hippocampus is connected with the anterior cingulate cortex and retrosplenial cortex (*4*, *16*).

In humans, evidence for structural differentiation between the anterior and posterior hippocamps is provided by age and Alzheimer’s Disease-related hippocampal volume reduction differences (*17*). Functional differences between the anterior and posterior hippocampus have also been proposed based on differential engagement during cognitive tasks (e.g., encoding vs. retrieval, emotion vs. cognition in anterior vs. posterior hippocampus, respectively) (*3*, *9*). Other fMRI research has suggested an anterior-posterior gradient in coarse-to-fine mnemonic spatiotemporal representations (*18*), such that anterior hippocampus supports more schematic or gist-like representations, while specific details associated with a given event are represented in posterior hippocampus (*6*, *8*). Similarly, other studies have suggested anterior-posterior hippocampal differences in pattern completion (i.e., integrating indirectly related events) and pattern separation (i.e., discriminating between separate but similar events) (*19*).

Resting-state functional connectivity (RSFC) fMRI studies in humans have provided additional insights into the hippocampal connectivity that underlies hippocampus-mediated cognition. RSFC exploits the phenomenon that even in the absence of any overt task, spatially separated but functionally related regions exhibit correlations in blood oxygen level-dependent (BOLD) signal (*20*–*24*). Group-averaged RSFC studies have found the hippocampus to be functionally connected to the default mode network (DMN) (*25*–*28*). The DMN, which is relatively deactivated by attention-demanding tasks, is thought to be important for self-referential processes, such as autobiographical memory, introspection, emotional processing and motivation (*26*). Other group-averaged RSFC studies have reported the anterior hippocampus to be preferentially functionally connected to anterior pieces of the DMN, while the posterior hippocampus was more strongly connected to the posterior DMN (*29*–*31*).

Recent precision functional mapping (PFM) studies have highlighted that RSFC group-averaging approaches blur individual-specific network boundaries and obscures fine-grained detail of network architecture in both the cortex and subcortical structures (*32*–*40*). The large amounts of RSFC data utilized (>300 minutes per subject) in PFM increase the signal-to-noise ratio (SNR) and allow for the detection of finer-grained functional neuroanatomical details in the cerebral cortex (*33*), cerebellum (*32*), basal ganglia, thalamus (*34*), and amygdala (*35*). In a small, deep-lying structure like the hippocampus, group-averaging RSFC data may be even more problematic.

The medial parietal cortex (mPC) is one of the main targets of hippocampal anatomical and functional connectivity (*4*, *16*, *41*–*45*), and was broadly considered part of the DMN (*25*– *28*). The mPC was defined as the swath of posterior midline neocortex between motor and visual regions that includes the retrosplenial cortex, posterior cingulate, and precuneus (BA 7, 23, 26, 29, 30, and 31). More recent studies revealed that parts of the mPC belong to the parietal memory network (PMN) and the contextual association network (CAN) (*23*, *33*, *46*, *47*). The PMN and CAN are immediately adjacent to the DMN in mPC, and therefore easily confounded. The identification of multiple different networks (DMN, PMN, CAN, FPN [fronto-parietal network]) in mPFC reflects the ongoing recognition of novel networks, subnetworks and organizational principles driven by PFM (*23*, *33*, *38*, *40*, *48*–*50*).

The DMN, PMN and CAN are all networks believed to be important for memory. The DMN and PMN have been associated with different aspects of episodic memory processing. Autobiographical retrieval, i.e., memory over a lifetime, preferentially increases activity in the DMN, whereas memory for recently experienced events preferentially engaged the PMN (*27*, *46*, *51*, *52*). During explicit memory tasks, activity within the PMN reduces in response to novel stimuli, but increases in response to familiar stimuli, where the degree of increased activity likely reflects attention to internal memory representations during retrieval (*46*, *53*). The CAN has been found to engage in contextual and visual scene processing, mediated in part by associations built from life experiences between objects or places and their scenes (*40*, *54*).

Here we utilized PFM to examine individual-specific, fine-grained, hippocampal-cortical connectivity in the Midnight Scan Club (MSC) data set (n=10 participants; 300 min. of rs-fMRI data/subject) (*33*). Due to the small size of the hippocampus, we also utilized additional highly-sampled, higher-resolution BOLD rs-fMRI data (2.6mm isotropic voxels; 2610 minutes; MSC06-Rep) from an independent dataset (*55*, *56*) as additional validation for the results. We generated individual-specific RSFC parcellations of the hippocampus, drawing on several advantages over group-averaging, including: (1) higher signal-to-noise ratio (SNR) in deeper subcortical structures without blurring individual differences in network features, and (2) more precise definition of individual-specific cortical functional network maps (i.e., DMN, PMN, CAN, FPN).

## RESULTS

To characterize the fine-grained functional connectivity of the hippocampus in individuals, we used a winner-take-all (WTA) approach to examine preferential connectivity to larger-scale cortical networks, such that each voxel was assigned to the cortical network with which it was most strongly functionally connected (see Methods; (*32*, *34*)). We utilized 15 individual-specific networks generated from the Infomap clustering algorithm (*33*), excluding two medial temporal lobe networks as they include the hippocampus (**Figure S1**).

### Anterior-Posterior Dichotomy in Hippocampus Functional Connectivity

Individual-specific WTA parcellations of the hippocampus (15 networks), revealed that the anterior hippocampus was most strongly connected with the default mode network (DMN) in all individuals (**Figure S2A)**. Half of the subjects also exhibited some connectivity of the anterior hippocampus to the contextual association network (CAN).

In all subjects the most posterior portion of the hippocampus (tail) was most strongly functionally connected to the PMN (**Figure S2A**). In MSC06, the lower resolution data (4 mm; 300 min.) showed the posterior hippocampus to be most strongly functionally connected to the fronto-parietal network (FPN). However, WTA of the higher-resolution data set (1.6 mm; 2610 min; MSC06-Rep), which supersedes the lower resolution data, also showed the posterior hippocampus to be most strongly connected to the PMN (**Figure S3**).

We quantified the proportion of the hippocampus preferentially connected to each cortical network in the WTA analysis (**Figure S2B**), which revealed that on average 56% of the hippocampus was most strongly connected to the DMN, 13% to the CAN, 14% to the PMN, and 2% to the FPN.

Given that the WTA parcellation scheme cannot account for more than one winning network within a voxel, we also considered the second strongest cortical connection for each voxel, following previously published procedures (*34*). We found that functional connectivity to the FPN was second to the PMN in the posterior hippocampus (**Figure S4**). Similarly, the runner-up to the DMN in the anterior portion of the hippocampus was always the contextual association network (CAN).

### Tail of Hippocampus Connected to PMN with Head/Body Connected to DMN

The WTA analyses using all 15 functional networks showed differences in network organization between anterior (DMN) and posterior hippocampus (PMN). To clarify this anterior-posterior dichotomy, we next utilized a two-alternative (DMN vs. PMN) forced-choice WTA approach (**Figure 1A**). In all 10 MSC subjects (**Figure 1B**) including higher-resolution validation data (MSC06-Rep; **Figure S3**), we found a separation between the anterior/middle (DMN) and most posterior (PMN) hippocampus. We also found that the DMN was most strongly connected to the anterior ~80% (62-88%) of the hippocampus on average, with the posterior ~20% (12-38%) of the hippocampus to the PMN.

**Figure 1.**
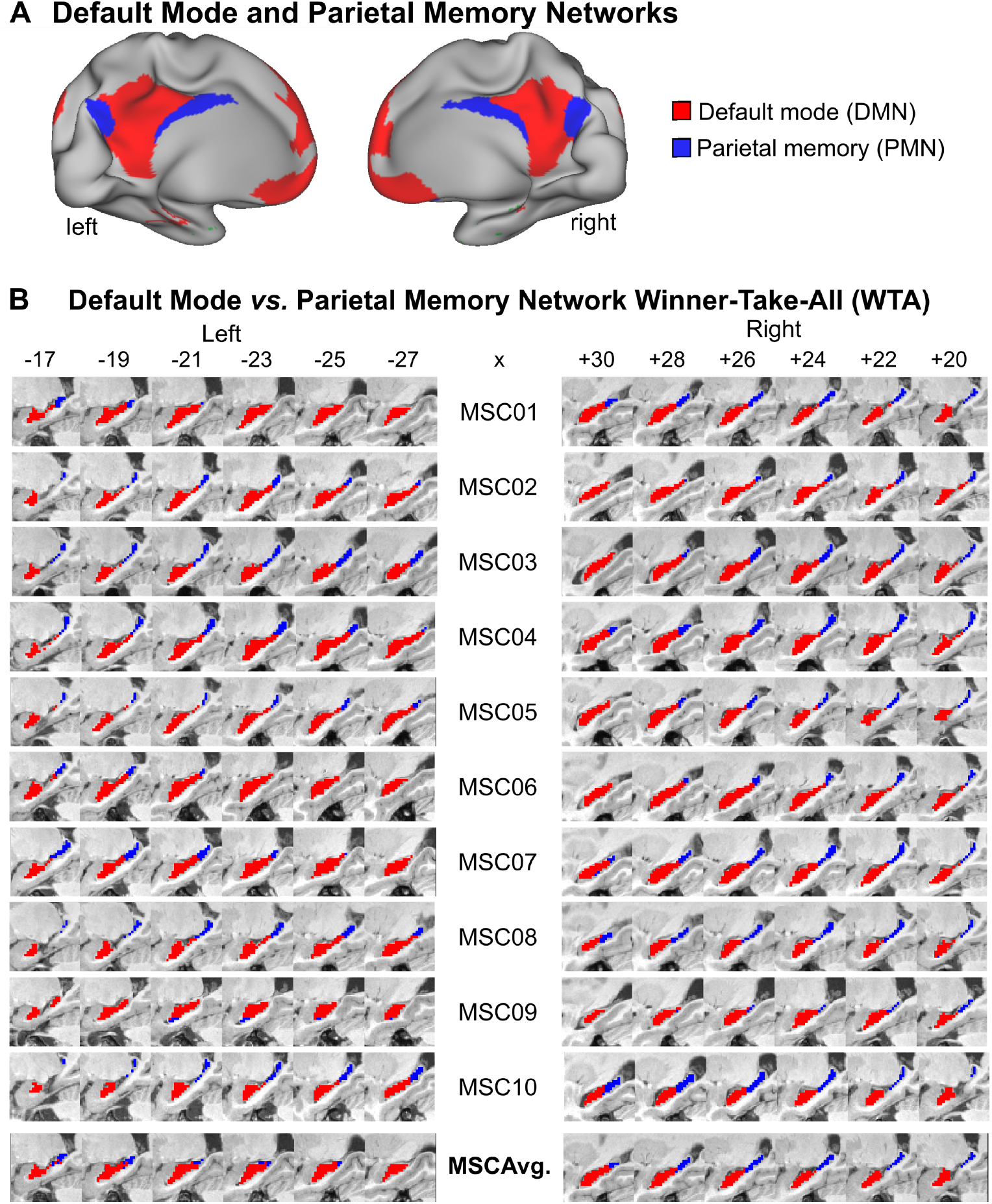
Hippocampal Parcellation Using a Two-network (Default [DMN] and Parietal Memory [PMN]) Winner-take-all Approach. **(A)** The default mode (DMN; red) and parietal memory (PMN; blue) networks are interdigitated in medial parietal cortex. **(B)** The DMN and PMN are organized along the anterior-posterior axis of the hippocampus such that the head/body portion is functionally connected to the DMN and the tail to the PMN.

Supplemental analyses demonstrated that the functional connectivity strength of each individual-specific hippocampal parcel to its winner cortical network was at least z(r) ≥ 0.2 (**Figure S5**, **Figure S6**). **Figure S7** visualizes the individual-specific DMN and PMN hippocampal parcels’ connectivity to the remaining cortical networks.

The differences in network connectivity between anterior and posterior hippocampus allowed us to generate subject-specific DMN and PMN hippocampal parcels (**Figure 1B**). Next, we needed to verify that these parcels could not have been generated by chance by showing that no other meaningful parcellation could be generated by this approach (**Figure 2**). We constructed participant-specific null distributions by conducting WTAs on the hippocampus using all possible pairs of networks (DMN, PMN, CAN, FPN excluded) and calculating the resulting parcels’ mean FC to the winner network. We found that the DMN and PMN parcels’ connectivity to their winning networks (DMN, PMN respectively) was significantly greater than for any other possible two-forced-choice WTA combination for each subject (p<0.001 for all comparisons for all subjects). The DMN parcels (left and right) were strongly positively correlated with DMN in every subject, but negatively correlated with the PMN (**Figure 2** top row). The PMN parcels were strongly positively correlated to the PMN, and uncorrelated with the DMN (**Figure 2** bottom row).

**Figure 2.**
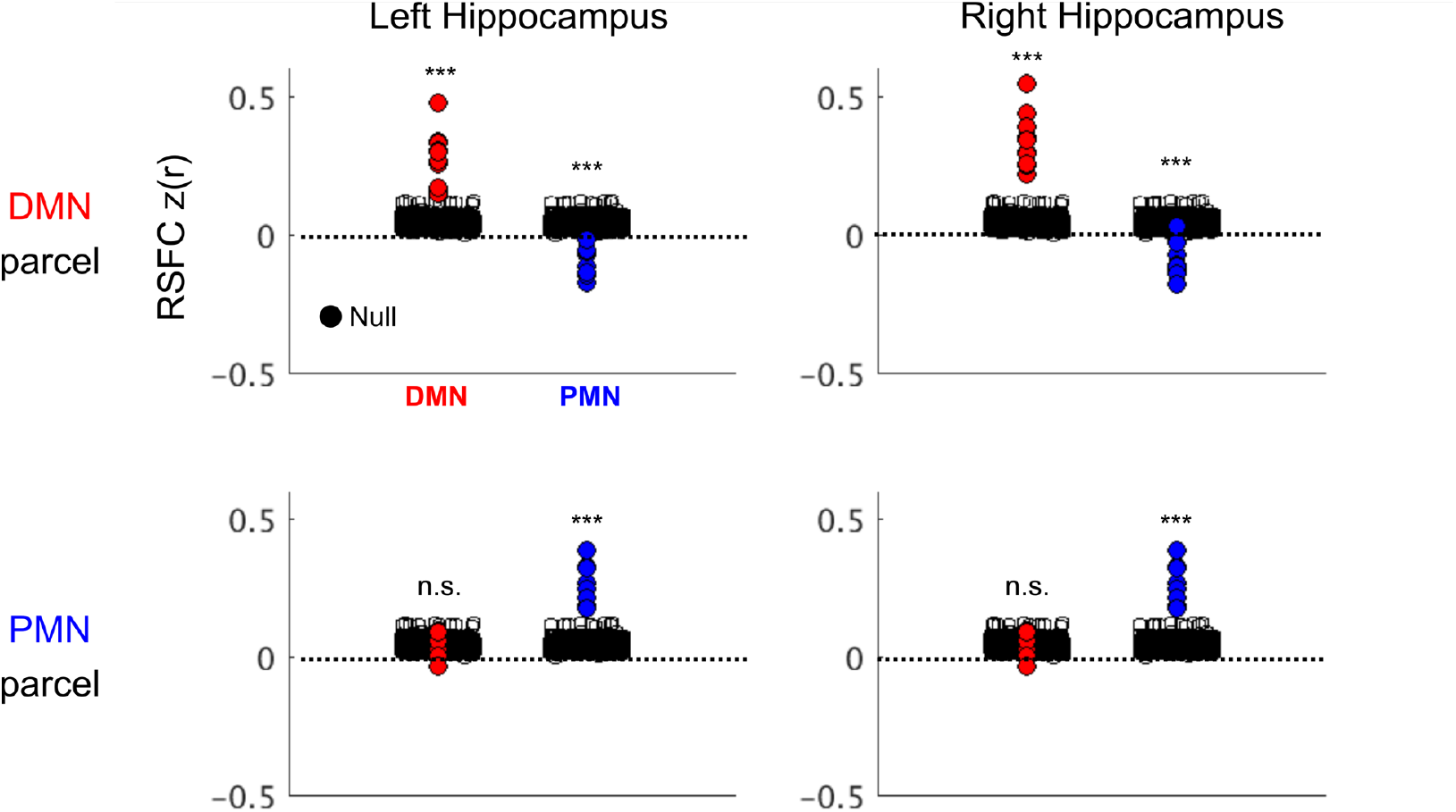
DMN and PMN Parcels’ Functional Connectivity to Cortical Networks. Displayed are the mean functional connectivity (FC) to the cortical DMN (red) and PMN (blue) for individual-specific WTA-derived hippocampal DMN and PMN parcels. Black circles indicate the null distribution, calculated from a hippocampal WTA between any pairs of networks and taking the generated parcels’ mean FC to its winner network. Although the null distribution for all participants are shown here, significance testing only occurred within-subjects against the participant-specific null distribution; *** p<0.001 for all subjects, n.s. p>0.05.

We then followed up the runner-up results (**Figure S4**; **Figure S7**), to similarly test whether the anterior and posterior hippocampus’s connectivity to cortical CAN and FPN respectively, were statistically significant. Replicating the above analyses, we computed the WTA on the hippocampus using two network candidates (CAN and FPN) (**Figure S8A).** We found that the RSFC of the resulting parcels was also significantly different from the null distribution (**Figure S8B**). We found that this CAN-FPN division strongly spatially overlapped with the DMN-PMN border (Dice = 0.4-0.9, median = 0.8).

### Connectivity of Hippocampal Parcels Matches Individual-Specific Network Boundaries

To visualize the cortical connectivity of the individual-specific hippocampal parcels (anterior, DMN; posterior, PMN), we displayed it over the previously defined (*33*) individual-specific cortical functional network boundaries (**Figure 3**; **Figure S9**). Using the hippocampal DMN and PMN parcels as regions of interests, we generated seed maps (**Figure 3** first two columns). Subtracting the individual-specific hippocampal seed maps (DMN parcel – PMN parcel) revealed sharp boundaries between the DMN and PMN in medial parietal cortex (**Figure 3** third column). Despite the DMN and PMN being immediately adjacent to one another in medial parietal cortex, differences in hippocampal-cortical connectivity between the anterior and posterior hippocampus re-traced the boundaries of their respective cortical networks (DMN, PMN) in an individual-specific manner. In contrast, the difference map between the group-averaged DMN- and PMN-parcel seed maps, did not show clear distinctions between the DMN and PMN networks (**Figure 3** bottom row).

**Figure 3.**
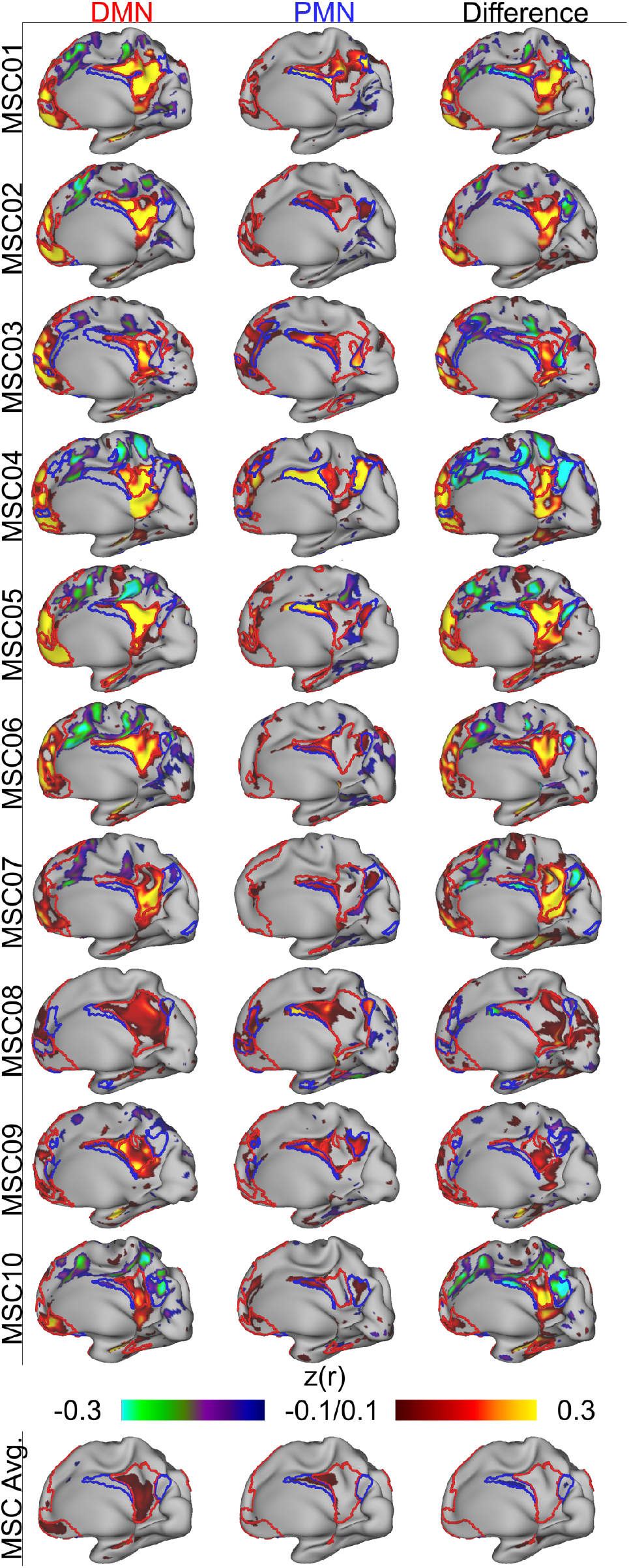
Functional Connectivity of Individual-Specific Hippocampal DMN and PMN parcels. The first two columns show the connectivity patterns of the anterior, default mode network (DMN) and the posterior, parietal memory network (PMN) parcels in the hippocampus. The third column depicts the difference maps of functional connections for the right hippocampus. The color scale for the last column is represented such that the warm colors represent greater DMN correlation and the cool colors represent greater PMN correlation. **Figure S9** shows the difference maps of the left hippocampal functional connectivity to the cortex.

### DMN-PMN Functional Connectivity Defines Functional Border in the Hippocampus

We tested whether the hippocampus’ functional connectivity was better explained by an anterior-posterior gradient or as modular network parcels (**Figure 4**). For each hippocampal voxel, we calculated the difference between its correlation with the DMN and the PMN (ΔFC = FC to DMN – FC to PMN). We tested whether hippocampal connectivity was better explained by a gradient or parcels using a one-way ANOVA (ΔFC ~ AP axis coordinates [gradient] or parcel identity). Across the group, both factors (r^2^_par_ vs. r^2^_grad_) explained roughly equal amounts of variance, with some inter-individual differences (**Figure 4**). The mean r^2^ across the 10 MSC subjects between parcels (mean r^2^_par_ = 0.48) and gradient (mean r^2^_grad_ = 0.46) were similar. Further, when replicating the gradient *vs.* parcel analysis in the more highly-sampled, higher spatial resolution MSC06-Rep data, we found that the parcel WTA identity also explained about the same amount of variance (r^2^_par_ = 0.77; r^2^_grad_ = 0.74) (**Figure S3D**). We also entered the gradient and parcel identity factors into a single ANCOVA to calculate the variance explained by one factor, after controlling for the other factor (**Table S1**). We found that both factors simultaneously explained separate variance, demonstrating both gradient and parcel organization in the hippocampus.

**Figure 4.**
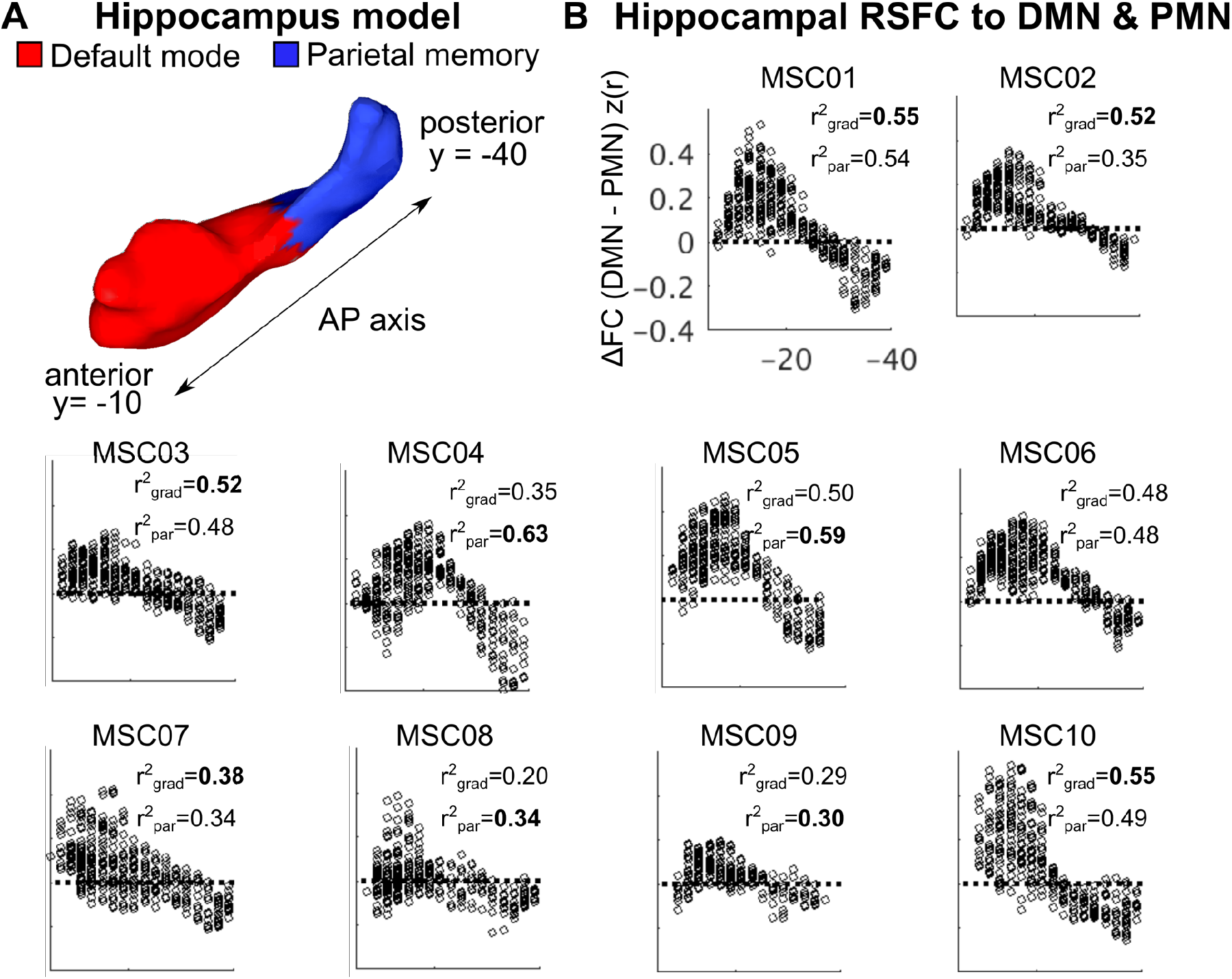
Hippocampal Functional Connectivity to Default Mode (DMN) and Parietal Memory Networks (PMN) along the Anterior-Posterior Axis. **(A)** Schematic of the hippocampus (MSC01) with the anterior-posterior (AP) axis drawn and the parcels outlined. For each subject, we determined each voxel’s position along the AP axis in the hippocampus as well as its parcel identity. **(B)** Scatterplots depicting the pairwise differences in functional connectivity (FC) to DMN and PMN as a function of coordinate position along the AP axis. The amount of variance in FC differences explained by the gradient (grad) and parcel (par) models are noted.

Having established that parcels explain just as much functional connectivity variance, we tested for a discernible functional border between hippocampal DMN and PMN parcels using receiver-operator-characteristic (ROC) analyses (**Figure S10**). We defined border voxels as adjacent to the other parcel within a 2-voxel radius and calculated the connectivity similarity (to cortex) for all possible pairs of border voxels. We sorted the similarity values based on whether the pair of voxels belonged to the same or different parcels (**Figure S10** histograms) and generated an ROC curve for each individual (**Figure S10**). We found that border voxels that belonged to the same hippocampal parcel (DMN or PMN) were significantly more similar than voxels that belonged to different parcels, for every individual (AUC=0.64-0.88; p<0.001). Thus, all 10 subjects had a discernible functional border between the DMN and PMN parcels.

### Anatomical Segmentation of Hippocampus Matches Functional Parcellation

WTA functional parcellation (**Figure 1**) segmented the hippocampus into DMN and PMN parcels. To test whether the functional parcellations revealed by the WTA approach mapped onto anatomical definitions of the hippocampal head/body and tail, we examined the spatial overlap between functional parcels and segments defined anatomically. We used anatomical landmarks (*57*) to select the coronal slice that demarcates the anatomical border between the body and tail of the hippocampus to create a partition separating the tail from the rest of the hippocampus (**Figure S11**). Specifically, we defined the end of the hippocampal body as the coronal slice where the fimbria fornix is posterior to the pulvinar nucleus of the thalamus (*57*). We found a high degree of spatial overlap between anatomical segments and WTA-defined parcels (Dice coefficient >0.7) across all subjects (**Figure S12**).

Next, we tested whether anatomically-defined hippocampal segments (head/body vs. tail) replicated the functional connectivity differences observed with functionally defined parcels (DMN, PMN; **Figure 3**). This validation analysis, using anatomically-defined regions of interest within the hippocampus, also found that functional connectivity patterns of the hippocampal head/body and tail differed starkly (**Figure 5**). That is, the head and body of the hippocampus were strongly preferentially connected to the DMN and the tail was strongly preferentially connected to the PMN.

**Figure 5.**
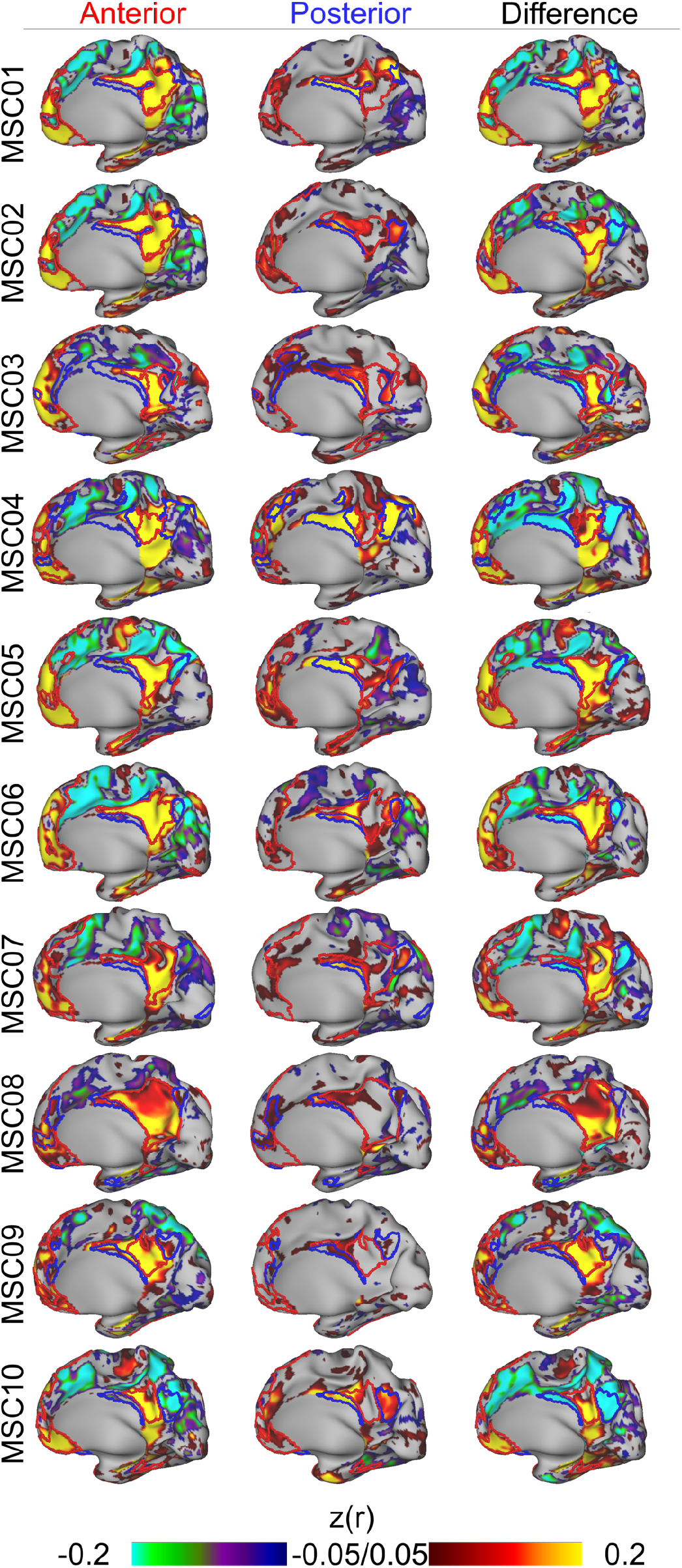
Functional Connectivity of Anatomically Defined Hippocampal Segments (Head/body vs. Tail). The hippocampus was split into two segments, head/body and tail based on anatomy. Functional connectivity seed map for head/body parcel (left), tail seed map (middle) and the difference between the two (right).

While the anatomical hippocampal segments mapped well onto the functionally-defined parcels, functional definitions are still advantageous because they do not require manual segmentation and seem to capture the true segmentation slightly better (**Figure S12**).

### Task-general Deactivations Specific to the DMN while Relatively Sparing the PMN

To validate the segregation of the DMN from the PMN, seen with RSFC and structural MRI, we next examined task-driven activations and deactivations during a mixed design, spatial coherence and noun-verb discrimination tasks. We found that robust task-driven decreases in activity were localized to the DMN, with less pronounced or no deactivations in the PMN (**Figure 6A**). The task-general decreases, the DMN’s defining characteristic, were significantly greater for the DMN than for the PMN networks in the cortex (**Figure 6B**; p<0.001), as well as the DMN and PMN parcels in the mPC (**Figure 6C**; p<0.001) and hippocampus (**Figure 6D**; p=0.02). Thus, the dichotomization into parallel DMN and PMN hippocampal-parietal circuits was borne out by anatomy, functional connectivity and task fMRI.

**Figure 6.**
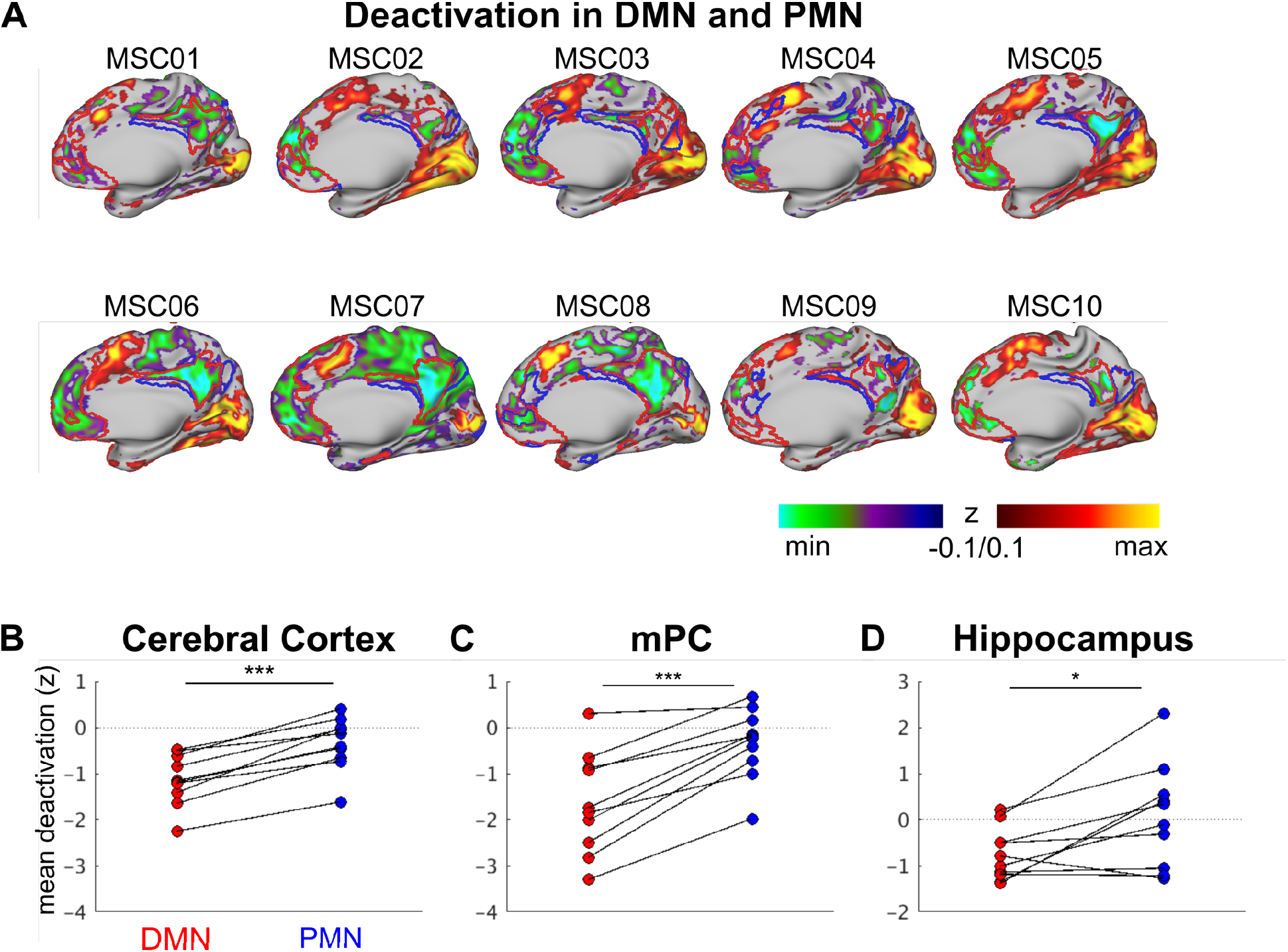
Task Deactivations in DMN and PMN. **(A)** Decreases in neural activity during task-based fMRI with functional connectivity defined network borders overlaid (DMN = red; PMN = blue). Mean task deactivations in DMN, PMN were calculated for **(B)** all of cerebral cortex, **(C)** the medial parietal cortex (mPC), and **(D)** the hippocampus. *** p<0.001; * p<0.05

## DISCUSSION

### Superimposition of Functional Parcels onto Hippocampal Gradients

Prior studies have largely conceptualized the organization of the hippocampus as a gradient—e.g., size of place field representations (*14*, *15*) or spatiotemporal scale of mnemonic representations (*6*, *8*). Using precision functional mapping (PFM) we documented overlapping gradient and parcel organization along the hippocampal anterior-posterior axis. We found the anterior 4/5^th^ of the hippocampus (head and body) to be preferentially functionally connected to the default mode network (DMN), with secondary connections to the contextual association network (CAN). In contrast, the posterior hippocampus (tail) was preferentially functionally connected to the parietal memory network (PMN), with secondary connections to the fronto-parietal network (FPN). The DMN, CAN, PMN and FPN are arranged as a topological ensemble across the hippocampus and the cortex, particularly along the midline in the mPC, such that these networks are immediately adjacent to one another.

To test whether these results were affected by data amount, quality or resolution, we used finer-grained (2.6mm) data from MSC06-Rep (2610 minutes), which again showed both parcellation and a gradient. The strong modularity of the hippocampus was validated by the localization of task-driven deactivations to DMN parcels, while relatively sparing the PMN.

**Figure 7.**
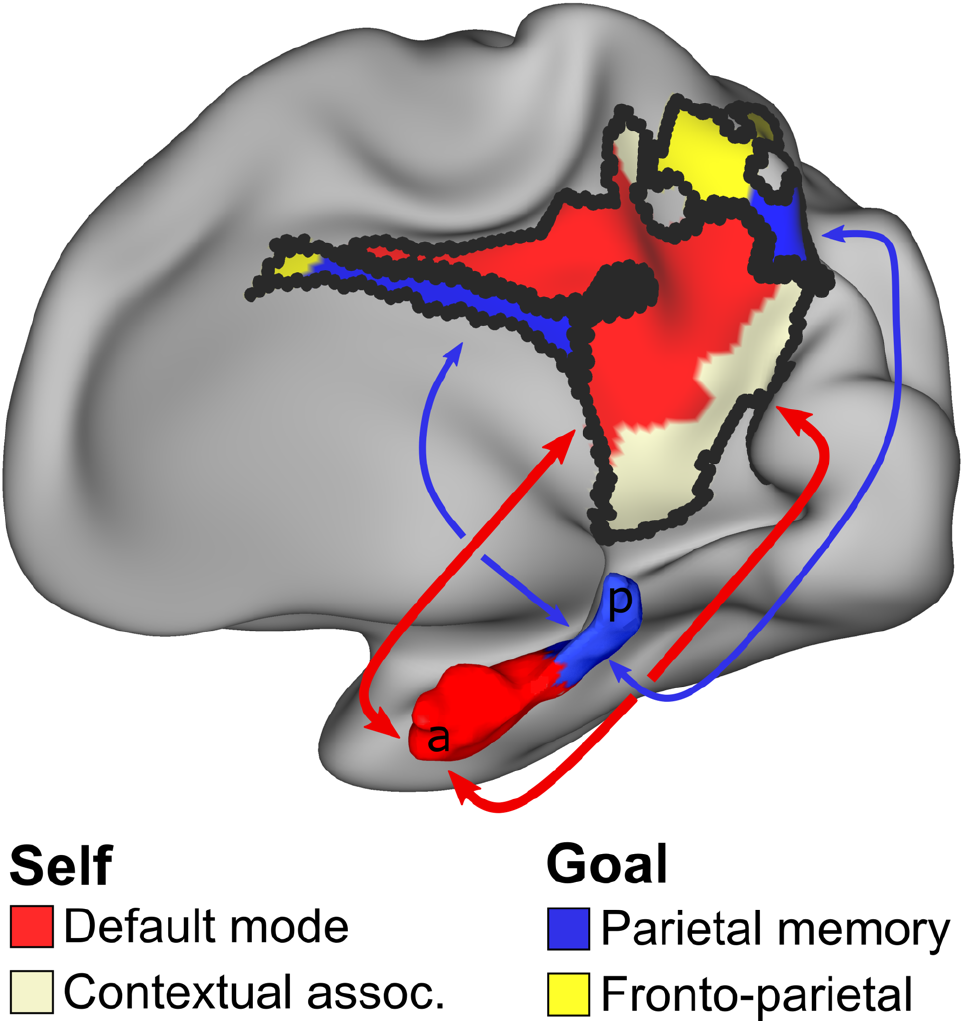
Schematic of Parallel Self- and Goal-Oriented Circuits between the Hippocampus and Medial Parietal Cortex. Medial parietal cortex is the primary target of hippocampal functional connectivity, but connectivity is segregated by functional network. The bulk of the hippocampus (anterior) is functionally connected to the default mode (DMN; red) and contextual association (CAN; pearl white) networks. The tail of the hippocampus is preferentially connected to the parietal memory (PMN; blue) and fronto-parietal networks (FPN; yellow). This functional connectivity dichotomy maps onto those parts of hippocampus and medial parietal cortex that deactivate during goal-oriented tasks (DMN, CAN) and those that do not (PMN, FPN). This functional organization suggests that human cognition can draw on two variants of hippocampal, medial parietal circuitry. The anterior circuit might support sequencing and navigating spacetime (*58*, *59*) in the service of the self, while the posterior circuit might carry out very similar operations in the service of accomplishing specific goal-directed, attention-demanding tasks.

### Anterior Hippocampus to Medial Parietal Cortex Circuit for Self-Oriented Processing

While the hippocampus, along with medial prefrontal cortex and posterior cingulate, have long been recognized as core regions of the DMN, our results more precisely define the anterior 4/5^th^ of the hippocampus as the hippocampal subregion that interacts with the cortical DMN, likely mediated by the parahippocampal gyrus (*60*). The DMN is thought to be important for autobiographical memory, emotion processing, prospective thinking, and social cognition (*25*, *26*). Thus, the DMN is generally thought to mediate introspective, self-oriented types of cognition (*61*).

We observed some intermixed functional connectivity to the CAN in the anterior hippocampus. In the higher-resolution (2.6mm isotropic voxels) and more highly-sampled (2610 minutes) data, we were able to replicate the observation that the anterior hippocampus is functionally connected to both the DMN and CAN. In fact, the CAN connectivity in the anterior hippocampus becomes more readily apparent with finer spatial resolution and larger amounts of data. In contrast to the functional border between head/body (DMN + CAN) and tail (PMN + FPN), the network representation of both DMN and CAN in the hippocampal head/body appears to be intermixed. The interdigitated nature of the DMN and CAN in the head/body of the hippocampus even with 2610 minutes of 2.6 mm data (MSC06-Rep) suggests that this is not caused by noise or blurring. This could be due to the fact that (i) the DMN proportionally occupies a larger part of the neocortex and/or (ii) the CAN is a subnetwork of the DMN (*38*, *40*, *48*, *62*).

Some have theorized that the CAN mediates the generation of predictions in top-down processing based on the learned associations between environmental features (e.g., between objects and their associated contexts) generated from a lifetime of repeating patterns of co-occurrences (*47*, *54*, *62*, *63*). Therefore, as contextual associations are important for episodic memory and spatial mapping, the anterior hippocampus (head and body) may be a zone of integration for both the DMN and CAN, playing a role in the circuity responsible for utilizing associative knowledge during episodic memory and affective processing. Others have argued that the hippocampus’ plays a role in binding item information within spatiotemporal contexts (*64*) and have suggested a role in scene processing (*65*).

The presence of both CAN and DMN connectivity may be highlighting the importance of integrating contextual, social, and affective information in the anterior hippocampus, which is consistent with prior notions of the anterior hippocampus’ interactions with the ventro-medial prefrontal cortex in schema generation (*6*, *9*, *66*). This integration is important for constructing and updating schemas, i.e., coherent worldviews of one’s environment in which the associated episodic memories, spatiotemporal contexts, social cognition and emotions are consistent with one another. Specifically, the anterior hippocampus-mPC circuit likely supports mental simulations based on autobiographical memory, theory of mind, self-referential judgements, etc. in order to guide expectations and comprehension of the interior life and external environment. A more general description of the anterior circuit’s function may be to support ordinal sequencing in spacetime (*58*, *59*) in the service of the self.

### Posterior Hippocampus to Medial Parietal Cortex Circuit for Goal-Oriented Processing

The circuit between posterior hippocampus, PMN, and medial parietal parts of the FPN may be part of a system that integrates PMN and FPN functions in order to allow attention-directed memory retrieval. The PMN is always deactivated relative to baseline by novel stimuli (*46*, *51*, *53*, *67*). The PMN is activated by familiar stimuli during explicit novel vs. familiar judgments, but fails to do so implicitly, without attentional focus on familiarity (*67*). The PMN’s task-driven activity patterns seem to reflect attention to relevant internal mnemonic representations (*46*). Meanwhile, the FPN supports executive functions (*22*), such as directing visual attention. This integration of PMN and FPN in the posterior hippocampus might involve (i) attention to relevant internal memory representations or schemas similar to the current environmental input (*46*), (ii) the retrieval of prior experiences that may be relevant for selecting task-appropriate responses (*10*, *68*), and (iii) selecting relevant novel information to update the appropriate internal memory representations (*69*, *70*).

While it is known that the hippocampus is involved in novelty-familiarity discrimination tasks (*69*, *71*), our results suggest that the tail (rather than the head/body) of the hippocampus may be more important for familiarity judgments of recently-seen stimuli (*3*) given the preferential connectivity of the tail to the PMN. The PMN may be crucial for long-term memory encoding by directing cognitive resources towards encoding novel information (*69*, *70*).

A notable finding in the present study is that the hippocampal tail was not dominantly connected to occipitotemporal cortex, but rather to medial parietal cortex. Prior studies have asserted that posterior hippocampus is functionally connected to visual/perceptual neocortical networks in support of fine, detailed, perceptually-rich representations of memories (*9*). We theorize that attention to and comparison with finer-grained mnemonic representations is necessary to determine whether the current environmental input is familiar or novel without necessitating preferential engagement of visual networks.

In addition to being strongly functionally connected to the PMN, the hippocampal tail was also connected to the FPN. Chen et al., 2017 found that episodic memory retrieval triggered by recently seen stimuli preferentially activated the PMN and parts of the FPN. This PMN-FPN interaction suggests that certain aspects of episodic memory retrieval, such as retrieval of relevant prior experience for task-appropriate responses, requires a broader set of network engagement. Further, engagement of the PMN and FPN during retrieval of task-relevant prior experience is consistent with the current finding that the tail of the hippocampus is preferentially connected to both the PMN and FPN.

### Comparisons between the Human Hippocampus and Animal Models

The basic organization of the hippocampus across rodents and primates is along the dorsal-ventral or posterior-anterior axis (*2*, *4*). However, in rodents, the majority of the hippocampus, i.e., dorsal and midtemporal third, is dedicated to processing visuospatial information while the neocortex is largely dedicated to processing somatosensory inputs and generating motor outputs (*58*). With the neocortical expansion in the primate brain to more complex, higher-order cognitive functions, more of the primate hippocampus is involved in processing this non-sensory information (*58*). Based on our functional parcellation of the human hippocampus, the more complex functions of the hippocampus, i.e., the self-oriented DMN parcel, is overrepresented compared to the rodent brain, consistent with the association cortex expansion in evolution. In comparison, the goal or task-oriented PMN hippocampal parcel, or the hippocampal tail, which is homologous to the visuospatial-oriented dorsal hippocampus in rodents, is relatively smaller. This hippocampal tail or rodent homologue exhibits connectivity to the medial parietal cortex (mPC), and reflects the conservation of anatomy and function in being involved in visuospatial processing in the mPC (*4*, *9*, *16*, *43*, *58*).

### Importance of Hippocampo-Neocortical Dialogue across Brain States

Prior group-averaged human functional connectivity studies, even those reporting connectivity differences between the anterior and posterior hippocampus, falsely presumed the hippocampus to be exclusively associated with the default mode network (DMN). The DMN was originally defined as the brain regions that collectively deactivated during goal-oriented, attention-demanding tasks, independent of the specific task demands (*61*). Subsequently, the DMN was also shown to be activated by a variety of self-referential, introspective tasks (*25*, *26*). The separation of brain regions into self-oriented (DMN) and task- or goal-oriented (not the DMN) is the first branch point when sorting brain regions according to their fMRI task activity profiles. This same basic branching is also apparent in RSFC data, where the earliest network branching is that between the self-oriented DMN and the task-oriented regions (*21*). Therefore, awake human hippocampal function has been primarily linked with the default mode.

The discovery that the tail of the hippocampus is specifically and selectively connected to task-oriented regions belonging to functional networks thought to be important for controlling attention and memory retrieval suggests that it is specialized for providing hippocampal computations to the task mode. Much larger parts of the hippocampus and cortex seem dedicated to the self or internally-directed default mode (*25*, *61*, *72*). Uncoupling of the retrosplenial cortex from other brain regions, including the hippocampus, is associated with disassociation-like behavior (*73*). The finding highlights the importance of the mPC for integrating environmental and sensory information with the egocentric perspective in self-oriented processes. Yet, it appears as if moment-to-moment goal-oriented activity is also dependent on the hippocampus. Differentiable, parallel loops between the hippocampus and corresponding mPC may be respectively specialized for supporting the self and action respectively. Thus, the mPC may be a bridge for this hippocampo-neocortical dialogue (*74*, *75*) for both self- and goal-oriented processes. This functional and anatomical segmentation of the hippocampus raises interesting questions about the effects of mode changes on hippocampal-neocortical interactions.

Analyses of high and low-frequency neural activity in rodents (sharp-wave ripples & theta) and humans (delta-band & infra-slow activity) show that the information flow encoded in high-frequency activity between the hippocampus and cortex reverses its direction during sleep, compared to the awake (resting) state (*76*–*79*), primarily in the DMN (*75*, *79*). It is theorized that the direction of low-frequency activity that coordinates this hippocampo-neocortical reciprocal dialogue reflects the cortico-hippocampal state changes between memory encoding (wake) and consolidation (sleep) (*79*–*81*).

Given the functional differentiation of the hippocampal tail from the head/body, does information flow during activity, rest and sleep, between the tail and neocortex follow that of the anterior hippocampus’ DMN parcel? Is functional connectivity to different neocortical targets (PMN, FPN) the main difference between the hippocampal tail and the remainder of the hippocampus while still performing the same general computation coordinated by theta oscillations (*58*, *82*), or does it perform slightly different computations via multiple theta generators (*83*)? Irrespective of the answers to such questions, it seems clear that hippocampal interactions with cortex are critically important across all brain states: sleep, wakeful rest and action.

## Funding

This work was supported by NIH grants NS115672 (A.Z.); NS088590, MH096773, MH124567, MH122066, MH121276 (N.U.F.D.); MH118370, AG13854 (C.G.); MH122389, MH109983 (C.S.); HD087011 (J.S.); NS110332 (D.J.N.); MH121518 (S.M.) and funds provided by the Kiwanis Neuroscience Research Foundation (N.U.F.D.), the Jacobs Foundation (N.U.F.D.), and the McDonnell Center for Systems Neuroscience at Washington University in St. Louis (A.Z., N.U.F.D., D.J.N).

## Author Contributions

Conceptualization, A.Z., D.F.M., E.M.G., N.U.F.D.; Methodology, software & formal analysis, A.Z., D.F.M., S.M., J.S.S., C.G.; Data curating & processing, S.M., T.O.L., A.N.V., D.A.; Writing – Original Draft, A.Z., N.U.F.D.; Writing – Review & Editing, A.Z., D.F.M., S.M., A.W.G., D.J.N. T.O.L., B.P.K., N.A.S., A.N.V., J.M.H., D.A., B.L.S., C.M.S., D.J.G., J.S.S., S.M.N., G.S.W., C.G., K.B.M., M.E.R., E.M.G., N.U.F.D.; Supervision, N.U.F.D.

## Competing interests

The authors declare no competing interest.

## Data availability

The Midnight Scan Club (MSC) dataset is publicly available (https://openneuro.org/datasets/ds000224 as is the MSC06-Rep data (https://openneuro.org/datasets/ds002766).

## MATERIALS & METHODS

### DATASET

#### Participants and Study Design

We employed the publicly available Midnight Scan Club (MSC) dataset for our analyses (https://openneuro.org/datasets/ds000224). Details of the dataset and processing pipeline have been previously described in (*33*). Here, we describe information about the data and methods that are pertinent to the current study.

The MSC consists of large quantities of fMRI data collected from 10 healthy, right-handed, adult participants (24-34 years; 5 females), who were recruited from the Washington University in St. Louis community. Participants completed 10 sessions of scanning that were 1.5 hour each, with all scan sessions completed within 7 weeks. Each session consisted of 30 minutes of resting-state fMRI (rs-fMRI), where subjects maintained open-eyes fixation on a white crosshair presented against a black background. The resting-state run was followed by fMRI scans for other tasks: a motor task, semantic judgement task, motion coherence task, and an incidental memory task.

The MSC06-Rep data is part of a publicly available dataset (https://openneuro.org/datasets/ds002766). Details of the dataset and processing pipeline have been previously described in (*55*).

#### MRI image acquisition

Participants were scanned on a Siemens TRIO 3T, beginning at midnight, across 12 sessions (2 sessions of structural MRI scans + 10 sessions of functional MRI scans). Structural images included four T1-weighted scans (TE = 3.74ms, TR = 2400ms, TI = 1000ms, flip angle = 8°, 0.8mm isotropic voxels, 224 sagittal slices), four T2-weighted images (TE = 479ms, TR = 3200ms, 0.8mm isotropic voxels, 224 sagittal, 224), four MRA and eight MRV scans, which were not used in the present study.

Functional images included 300 minutes of eyes-open resting-state fMRI BOLD data (10 sessions x 30min/session) and 350 minutes total of task fMRI BOLD data using a gradient-echo EPI BOLD sequence (TE = 27ms, TR = 2.2s, flip angle = 90°, 4mm isotropic voxels, 36 axial slices). Gradient echo field map images (one per session) were acquired with the same parameters. See Gordon et al. (2017) for more details (*33*).

One participant (MSC06) underwent an additional 87 imaging sessions on a Siemens Prisma 3T MRI scanner, consisting of fMRI scans with higher resolution (gradient-echo EPI BOLD sequence: multiband factor 4, TE = 33ms, TR = 1.1 s, flip angle = 84°, 2.6mm isotropic voxels, 56 axial slices). These additional resting-state fMRI scans were conducted as part of another study independent of the MSC data collection (*55*), but are labeled as MSC06-Rep in the present paper. See (*55*) for more details on image acquisition parameters and procedures. We used MSC06-Rep as a means of not only validating the results from the original 10 MSC subjects, but also as a way to address potential concerns from the MSC data with regards to lower spatial resolution, partial voluming effects and signal contamination from immediately adjacent grey matter in the MSC dataset.

### STRUCTURAL & FUNCTIONAL MRI DATA PROCESSING

The preprocessing stream, individual-specific cortical surface generation, mapping of BOLD data to individual-specific cortical surfaces, and Infomap-derived individual-specific for the MSC rs-fMRI data have all been previously described in greater detail in Gordon et al., (2017) and other papers (*32*, *33*, *84*). Here, we briefly describe the steps below.

#### Structural MRI

Cortical surfaces were generated according to procedures described in Laumann et al. (2015) (*39*). Each participant’s averaged T1-weighted image was run through FreeSurfer v5.3’s recon-all processing pipeline to create the anatomical surface, which was double-checked and manually edited using Freeviewer to ensure accuracy. Surfaces were then registered into fs_LR_32k surface space with the Multi-modal Surface Matching algorithm described in Glasser et al. (2016) (*85*).

#### Functional MRI (fMRI) preprocessing

All fMRI data were preprocessing in volume space to maximize cross-session registration, which involved slice-time correction, intensity normalization, and within-run head motion correction. The functional MRI data were registered to Talairach atlas space using the subject-specific averaged T2-weighted and T1-weighted structural image. We used the mean field map to apply a distortion correction for each participant before resampling into 2mm isotropic resolution. These steps were combined into a single interpolation using FSL’s applywarp tool (*86*).

#### Resting-state fMRI (rs-fMRI) data preprocessing

We further preprocessed the rs-fMRI data to reduce spurious effects that are likely to be unrelated to neural activity using a motion censoring procedure described in Power et al. (2014) (*87*). The motion censoring procedure in combination with our other preprocessing steps have been demonstrated to be the current best practices in the field for reducing motion artifacts (*88*). Motion-contaminated frames were identified based on a framewise displacement (FD) > 0.2mm or a temporal derivative of the root mean squared variance over voxels (DVARS) > 5.36. Two subjects (MSC03, MSC10) required additional correction for artifactual high-frequency motion in the phase encoding direction (anterior-posterior) as previously described (*33*, *84*).

After motion censoring, the data was further preprocessed with the following additional steps: (1) demeaning and detrending, (2) interpolating censored frames with least-squares spectral estimation, (3) temporal band-pass filtering (0.005 Hz < f < 0.01 Hz), and (4) multiple regression of nuisance variables, which include the global signal, principle components of ventricular and white matter signals (described below in ‘‘Component-based nuisance regression’’), and motion estimates derived from the Volterra expansion (*89*), applied in a single step to the filtered, interpolated BOLD time series. Finally, censored volumes were removed from the data for all subsequent analyses. Application of the temporal masks resulted in retention of 5704 ± 1548 volumes per subject (range of 2691-7530) or ~209 ± 57 min. Thus, even the subject with the least amount of data after motion censoring still retained around 100 minutes of rs-fMRI data.

For MSC06’s additional 2.6mm resolution data, the data were processed in the same manner described above. However, (1) FD measurements were corrected for artifactual high-frequency motion in the phase encoding direction, (2) the FD threshold for motion censoring was 0.1mm, and (3) the DVARS threshold for motion censoring was 6.

The cortical data were then registered to the surface (see above “Structural MRI”). The cortical surface data and volumetric subcortical and cerebellar data were combined into CIFTI data format using the Connectome Workbench toolbox (*90*). Voxels in the cerebellum and the subcortical structures (which include the hippocampus, thalamus, caudate, putamen, pallidum, nucleus accumbens, and amygdala) were derived from the FreeSurfer segmentation of each subject’s native averaged T1-weighted image and manually edited by expert neuroanatomists to ensure utmost accuracy in grey matter segmentation. These were then transformed into Talairach atlas space. Finally, the cortical surface functional data were smoothed (2D geodesic, Gaussian kernel, σ = 2.55mm). Due to the small size of the hippocampus along with other subcortical structures, we did not spatially smooth data within the volume and we up-sampled the fully processed data to 2mm isotropic voxels.

#### Component-based nuisance regression

The temporally filtered BOLD time series underwent a component-based nuisance regression approach followed in several other papers (*32*, *34*, *91*). We built nuisance regressors based off of individual-specific white matter and ventricle masks, which were segmented by FreeSurfer (*92*), spatially resampled, and registered to the fMRI data. Due to the fact that voxels at the edges of the brain are highly susceptible to motion and CSF artifacts (*93*, *94*), we created another nuisance mask specifically for the extra-axial compartment by thresholding the temporal standard deviation image (SD_t_>2.5%), excluding a dilated whole brain mask (*95*, *96*).

We applied dimensionality reduction to the voxel-wise nuisance time series as outlined in CompCor (*96*). However, the number of retained regressors was determined for each noise compartment by orthogonalizing the covariance matrix and retaining components ordered by decreasing eigenvalue up to a condition number of 30 (λ_max_/ λ_min_ > 30). The columns of the design matrix *X* comprised the retained components across all compartments, the global signal and its first derivative, and the six motion correction time series. Since the columns of the design matrix *X* may exhibit collinearity, we applied a second level SVD to *XX*^T^ to overcome potential rank-deficiency in the design matrix. This imposed an upper limit of 250 on the condition number. The regressors were applied in a single step to the filtered, interpolated BOLD time series.

#### Distance-based regression of adjacent grey matter cortical tissue

The hippocampus is in close proximity to some cortical areas (e.g., the medial temporal lobe), which results in signal contamination and spurious correlations between hippocampal voxels and adjacent grey matter vertices. To mitigate these effects, we regressed out the BOLD time courses of adjacent cortical grey matter tissue that were within 20mm of a given hippocampal voxel to remove this spurious functional connectivity for each voxel. We quantified the Euclidean distance between every hippocampal voxel and every grey matter vertex in order to determine the tissue signal that needed to be regressed out. We then took the average time course of these adjacent vertices and regressed it out of hippocampal voxels. This follows similar strategies taken in previous work on subcortical functional connectivity (*32*, *34*, *97*).

### METHODS & STATISTICAL ANALYSIS OF HIPPOCAMPUS FUNCTIONAL CONNECTIVITY

#### Manual tracing of the hippocampus

T1-weighted MRI data initially underwent automated segmentation using Freesurfer version 5.3, followed by manual editing of hippocampal results using ITK-SNAP software by a single highly-experienced rater (D.A.). For this procedure, outlines were inspected and adjusted in the coronal view of the T1-weighted image from posterior to anterior sections. The segmentations were subsequently modified in the axial and sagittal views. The left and right hemispheres were independently outlined. Anatomical boundaries generally followed the approaches of Watson and Thompson, with reference to an anatomic atlas (Duvernoy, 2005; Thompson et al., 2011; Watson et al., 1992).

#### Infomap clustering of cortical resting state networks

All sessions were concatenated together for each individual before proceeding with the Infomap community detection algorithm for individualized cortical resting state network mapping. Due to the individual variability of cortical network organization as shown in previous studies (*33*, *98*), the current study’s precise characterization of hippocampal organization requires that cortical networks are defined within individuals. Using the Infomap algorithm for community detection, we were able to define 17 cortical networks for each individual MSC subject as previously published (*32*) (**Figure S1**). Two medial temporal lobe networks were excluded due to the distance exclusion criterion and poor signal-to-noise ratio present in these regions. These individually-defined cortical resting-state networks were then used to conduct a winner-take-all (WTA) parcellation of the hippocampus described below.

#### Winner-take-all (WTA) parcellation of the hippocampus

We followed previously established WTA approaches for functional parcellation of non-neocortical structures (*32*, *34*) and applied it to the hippocampus. For each subject, the distance regressed, BOLD time courses for all 10 sessions were concatenated together for the entire brain. For each given hippocampal voxel, we calculated the average BOLD time course of all cortical vertices greater than 20mm away from the hippocampus that made up a particular cortical network for all 15 networks. The correlation between every cortical network and the hippocampal voxel were calculated, where the cortical functional network with the greatest, positive correlation strength was declared the winner in the hippocampal voxel.

Runner-ups or the second-place winners of the WTA were identified following previously established procedures (*32*, *34*). In short, we considered voxels as having multiple network connectivity, i.e., having a runner-up network, if their correlation strength to the second-place winner was at least 66% of the winning correlation strength.

#### Generation of binary masks for the default mode and parietal memory networks

We re-ran the WTA analysis described above on the hippocampus using just the default mode (DMN) and parietal memory (PMN) networks as potential winner networks to create masks for these two networks, splitting the hippocampus into two parcels for each MSC individual. All subsequent analyses relied on this individualized DMN-PMN WTA parcellation.

#### Hippocampal-cortical functional connectivity maps

The resulting parcels generated from the WTA approach were used to calculate the functional correlation to all cortical vertices. The mean time course for a parcel was calculated before correlating it with every vertex on the cortex with correlation strengths Fisher z-transformed. The resulting functional connectivity maps were plotted with individual Infomap-generated cortical network boundaries overlaid on top.

#### Anatomical segmentation of the hippocampus

The head and body vs. the tail of the hippocampus were anatomically defined by landmarks identified in Daugherty et al., (2015) (*57*). The landmarks were identified in the session-averaged T1-weighted structural image for each individual to identify the coronal slice in which the fornix appears posteriorly to the thalamus (*57*). Identification of the coronal slice was double-checked for accuracy by a neuroradiologist (J.S.S.). All hippocampal voxels posterior to said coronal slice was considered the tail of the hippocampus whereas all voxels anteriorly were considered the head/body of the hippocampus. The subsequent anatomical segmentation was then used to calculate the mean time course for each anatomical segmentation and connected to the cortical vertices. These correlation strengths were then Fisher z-transformed before being plotted with individual cortical network boundaries.

#### Task Deactivations

Task fMRI data were processed as previously described (*33*, *98*). We used a pair of mixed block/event-related design tasks which comprised language and perceptual task trials in order to model task-based deactivations that are traditionally associated with the default mode network. The mixed design task began with a task cue followed by a block of jittered trials in each task, modeled after (*99*). The ‘‘language’’ task trials consisted of single words where participants determined whether the words were nouns or verbs. The ‘‘perceptual’’ task trials consisted of Glass dot patterns (Glass, 1969) that were either at 50% or 0% coherent. Participants had to determine whether the dots were arranged concentrically, i.e., were the dots coherent?

After standard fMRI preprocessing, task fMRI data were entered in a General Linear Model (GLM) separately for each session from each individual using in-house IDL software (FIDL) (*100*). The mixed design tasks were modeled jointly in a single GLM with separate event regressors for onset and offset cues from each task, trials in each task (nouns and verbs for the language task, 0% and 50% for the perceptual task), and a sustained block regressor for the task period. Event regressors were modeled using a finite impulse response approach consisting of delta functions at each of 8 time points, allowing for the more complete modeling of different HRF shapes (*101*). Deactivations were identified using a contrast of the third and fourth time points from all conditions in the mixed design tasks (against an implicit, unmodeled, baseline).

## Supplementary Materials

**Figures**

**Figure S1.**
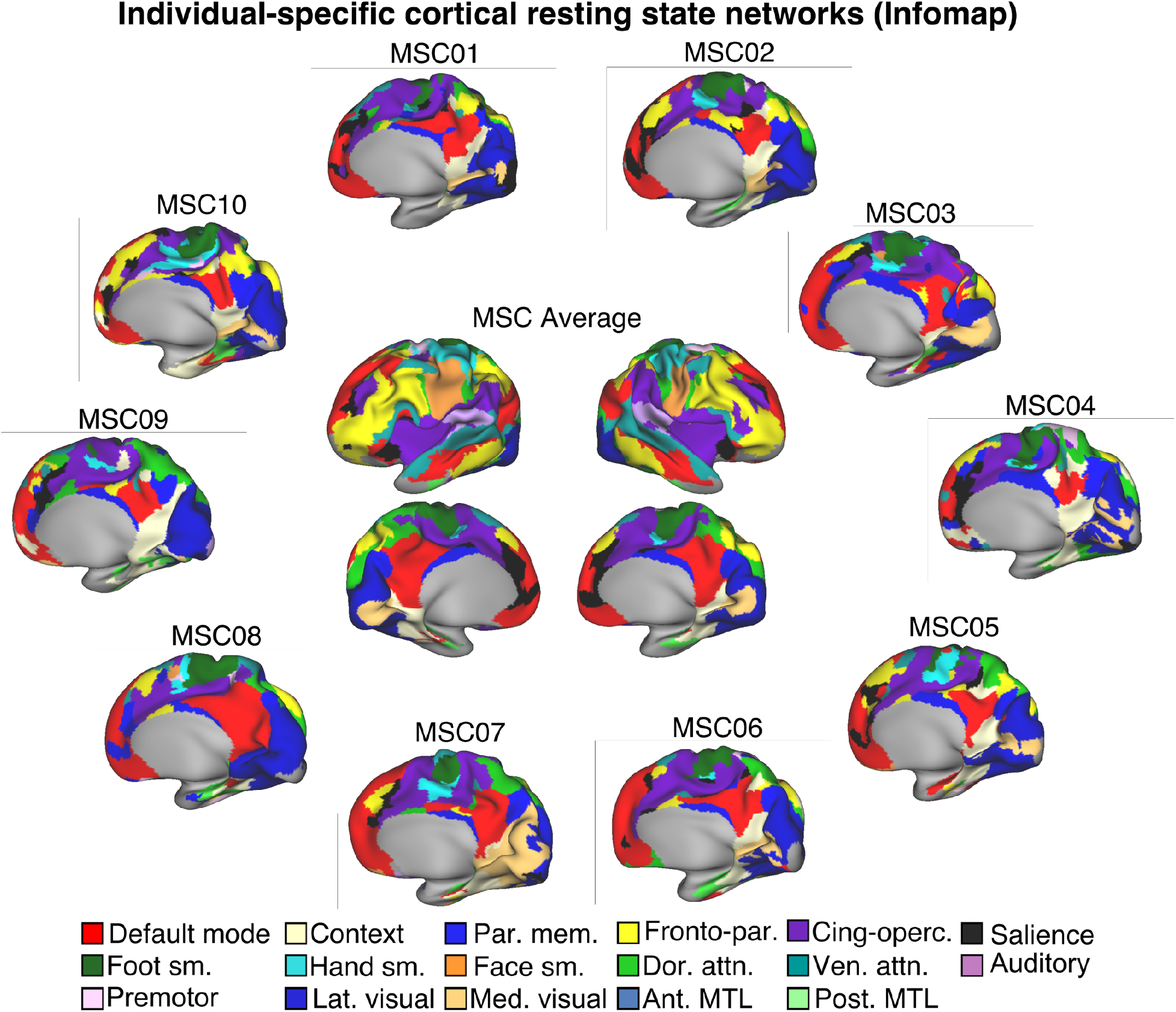
Individual-specific cortical resting state networks. Using the Infomap community detection algorithm, cortical resting state networks were defined for each individual MSC subject as well as the MSC Average. These individually-specified cortical networks were then used for the winner-take-all (WTA) parcellation of the hippocampus.

**Figure S2.**
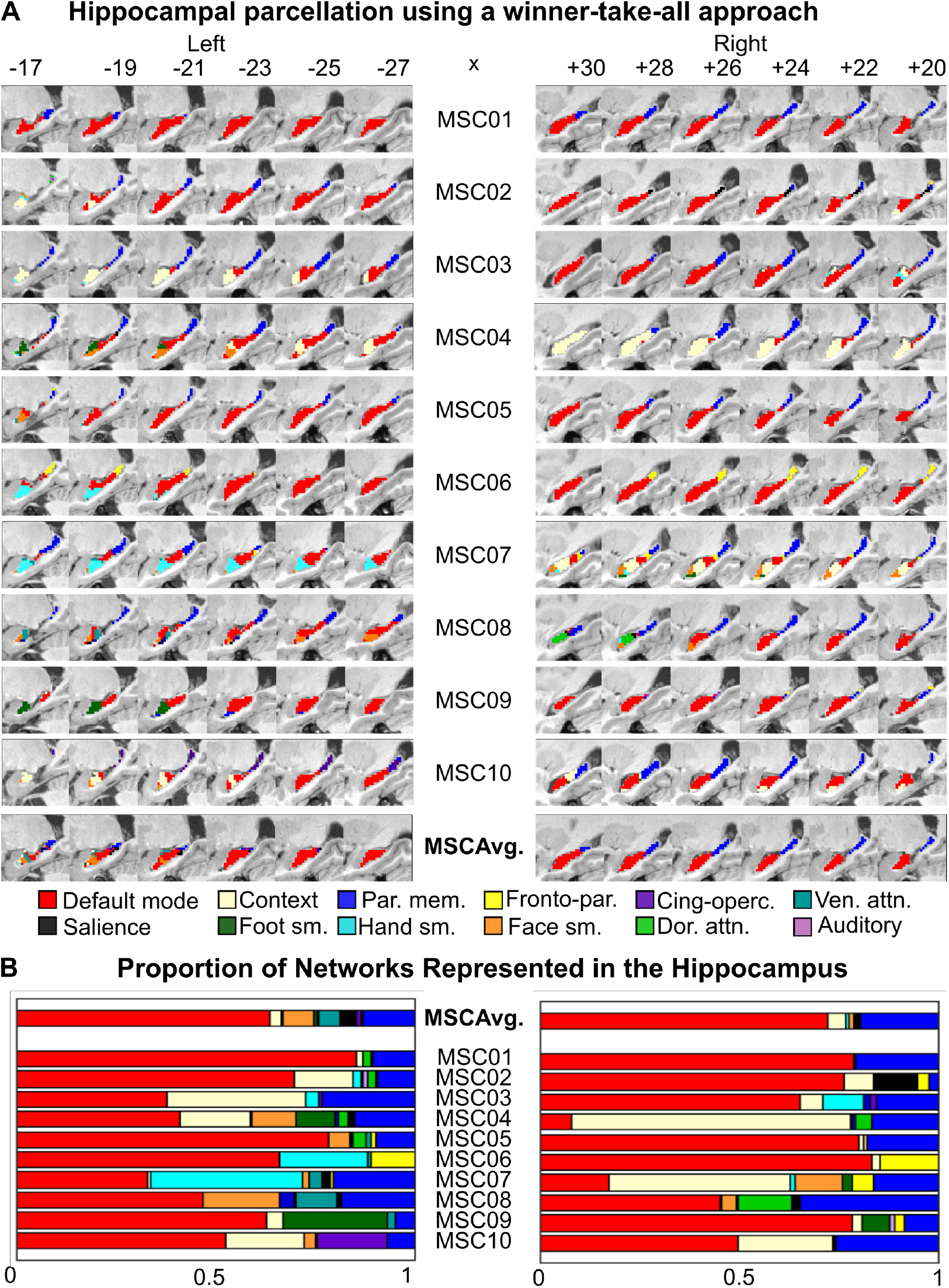
Hippocampal Parcellation Using a Winner-take-all Approach (WTA). **(A)** Anterior-posterior dichotomy in default mode network (DMN) and parietal memory network (PMN) representations in the left and right hippocampus for the MSC Average and each subject. Sagittal slices shown. **(B)** Quantification of the proportion of networks represented in the left and right hippocampus for each subject and MSC Average. High DMN (red) correlations occupied the largest proportion of hippocampal voxels and were primarily located in the anterior regions, while high PMN (blue) correlations primarily occupied the posterior hippocampus.

**Figure S3.**
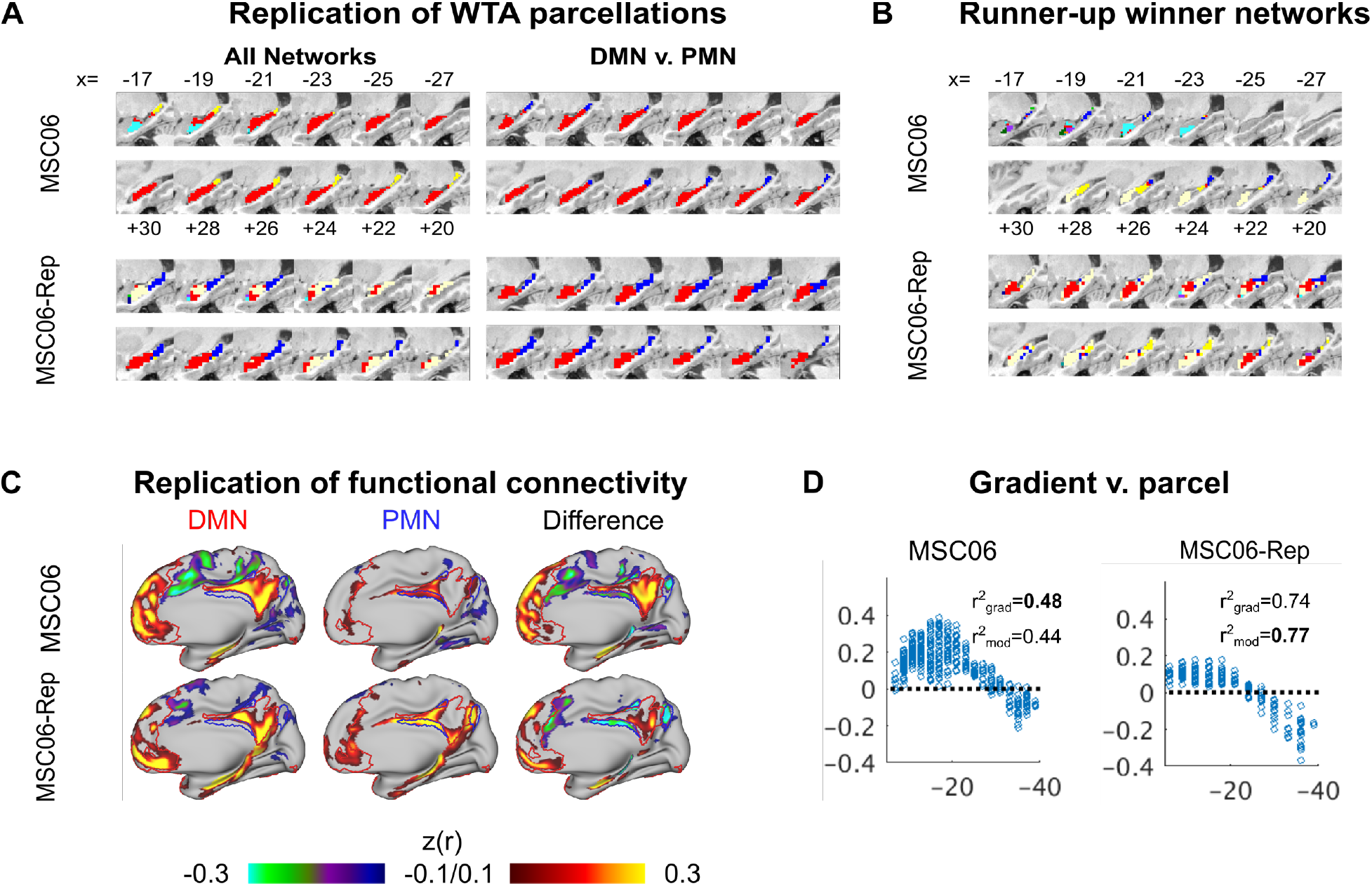
Replication of WTA parcellation in higher-resolution, higher-sampled BOLD rs-fMRI data from MSC06. Top panel shows the original network localization in the hippocampus for MSC06 using all potential winner networks and the bottom panel shows a replication of MSC06’s WTA parcellation using higher resolution data (2.6mm isotropic voxels).

**Figure S4.**
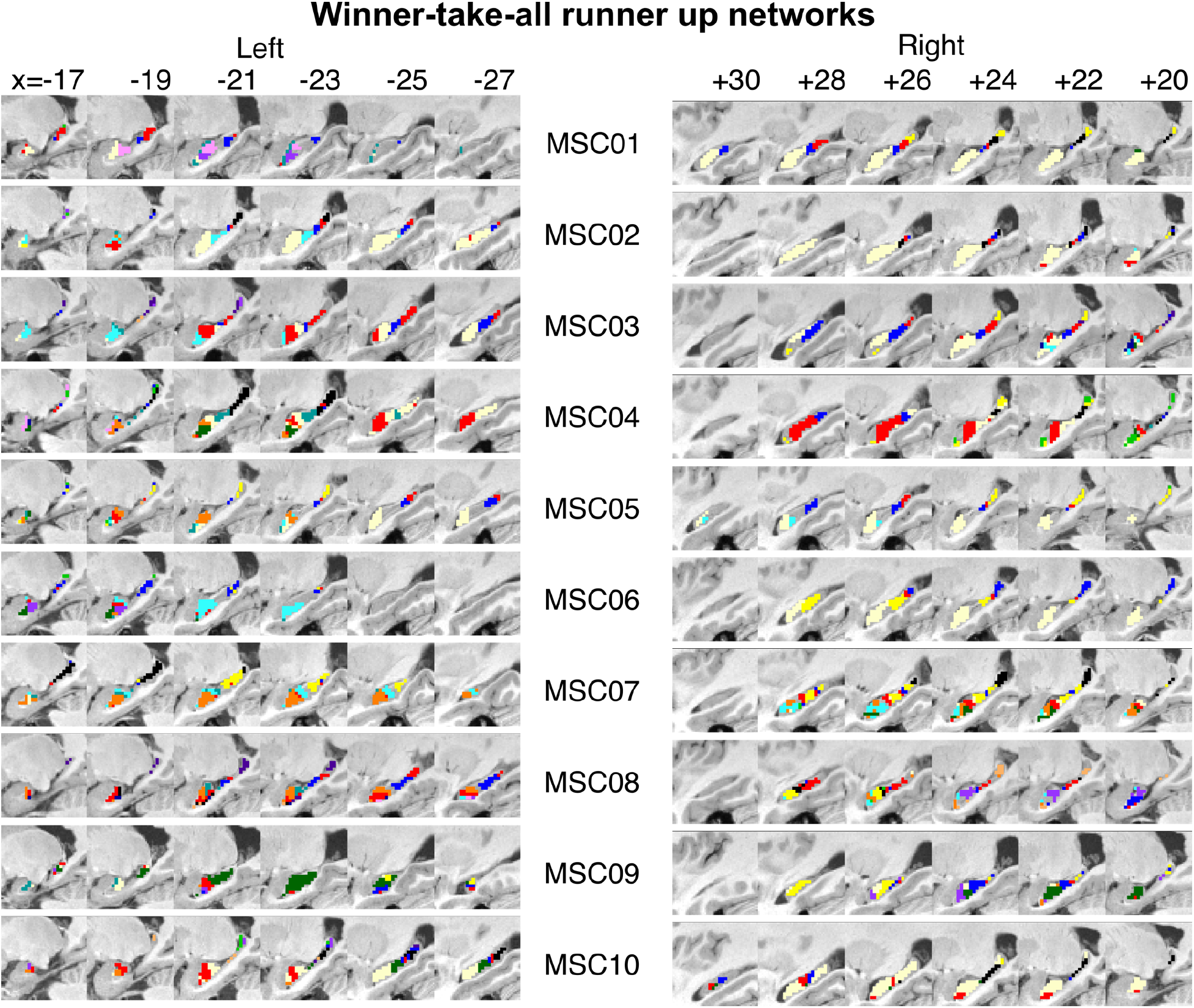
Runner-up winner take all (WTA) parcellation of the hippocampus. Pictured are the runner-up or second-place winning networks of the WTA parcellation of the hippocampus for all 10 MSC subjects. Not all hippocampal voxels were categorized as having multiple network connectivity; hence, some voxels are not colored. Voxels that were categorized as having runner-up network connectivity was based on if the runner-up correlation strength was at least 66.7% of the winning correlation following previously-established procedures (*32*, *34*).

**Figure S5.**
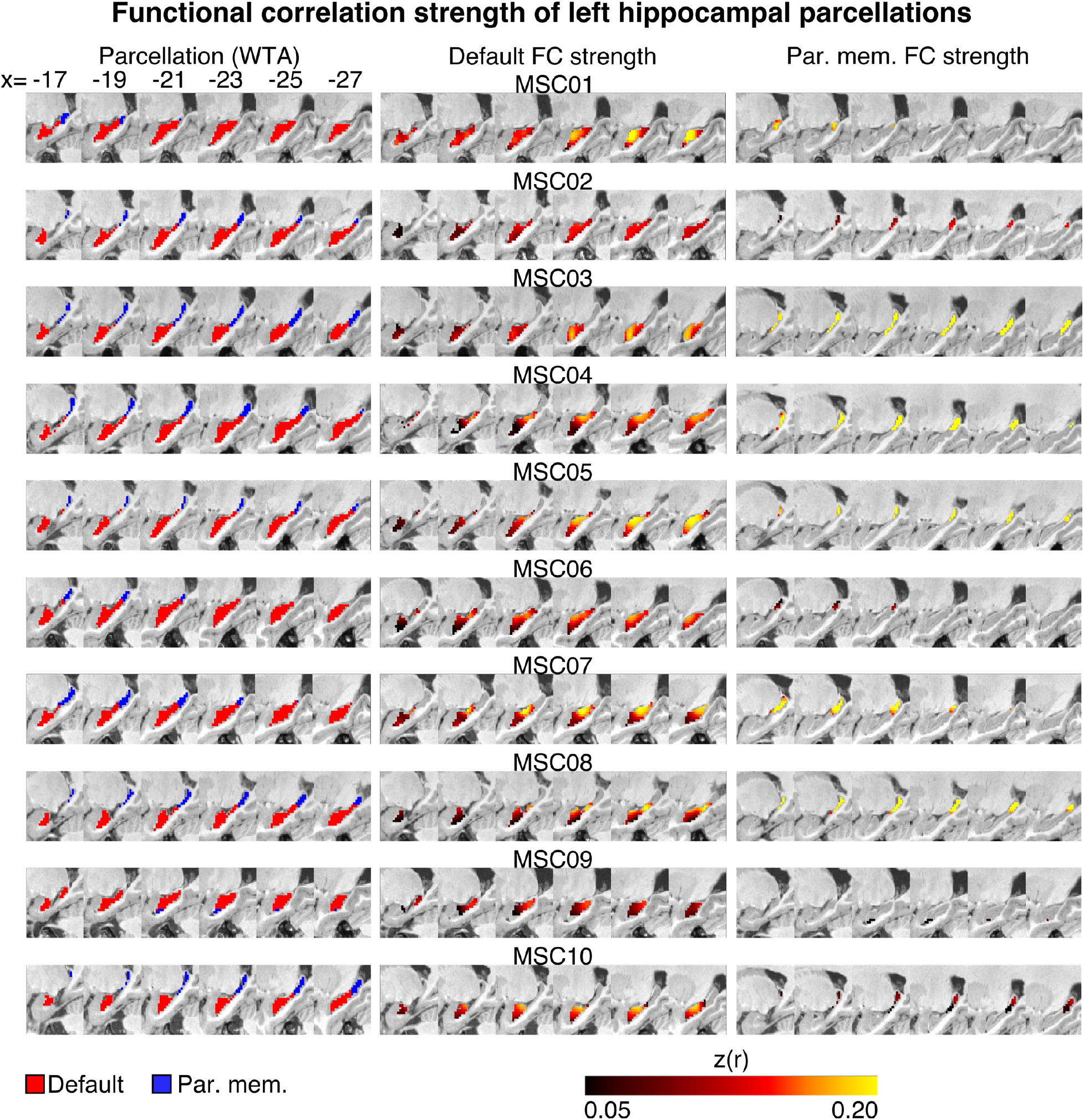
Functional correlation strength of WTA parcellations in the left hippocampus. The 2-network (DMN vs. PMN) WTA parcellation (left) of the left hippocampus is shown side by side with the DMN (center) and PMN (right) correlation strength to their respective winner networks.

**Figure S6.**
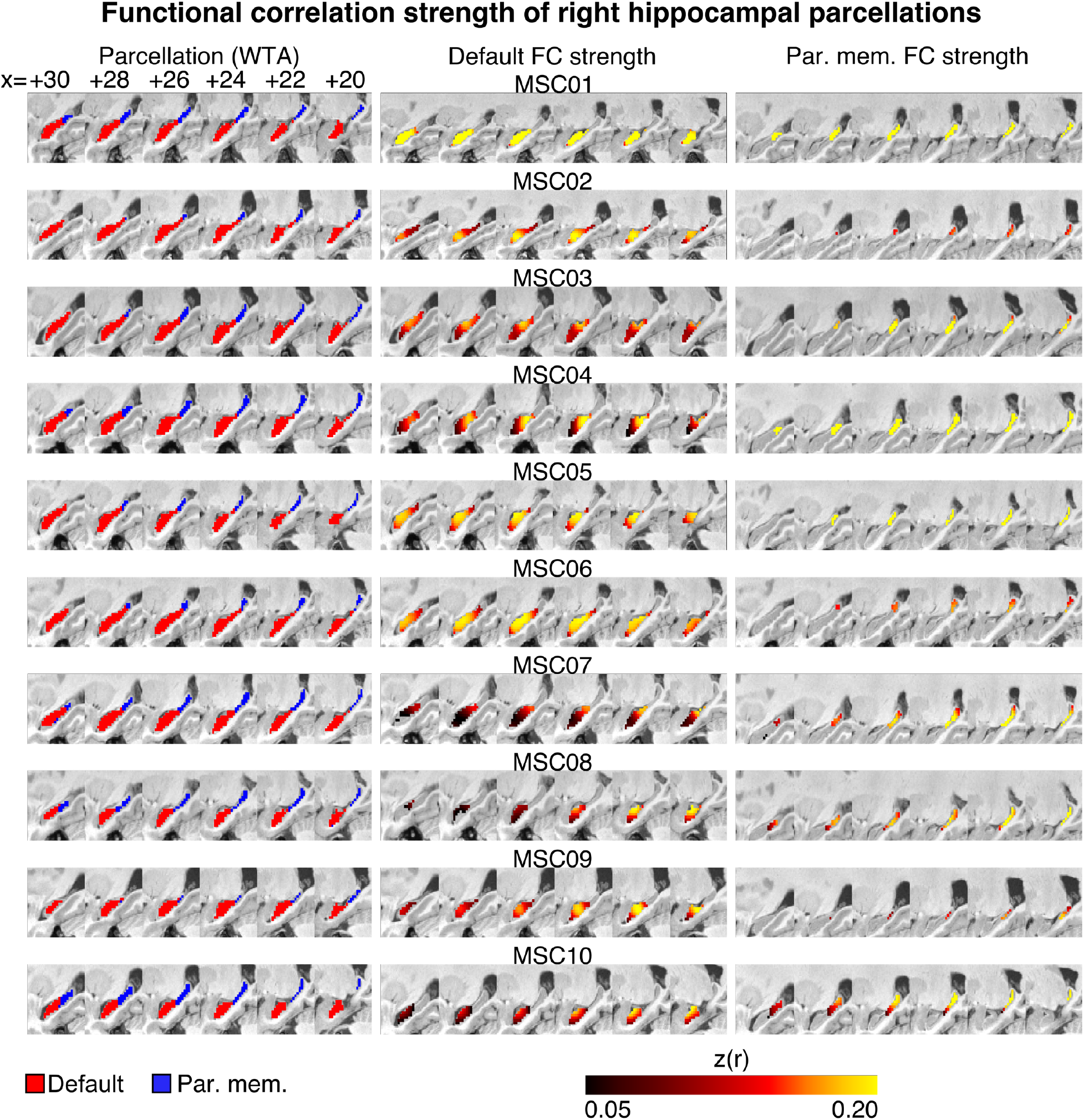
Functional correlation strength of WTA parcellations in the right hippocampus. The 2-network (DMN vs. PMN) WTA parcellation (left) of the right hippocampus is shown side by side with the DMN (center) and PMN (right) correlation strength to their respective winner networks.

**Figure S7.**
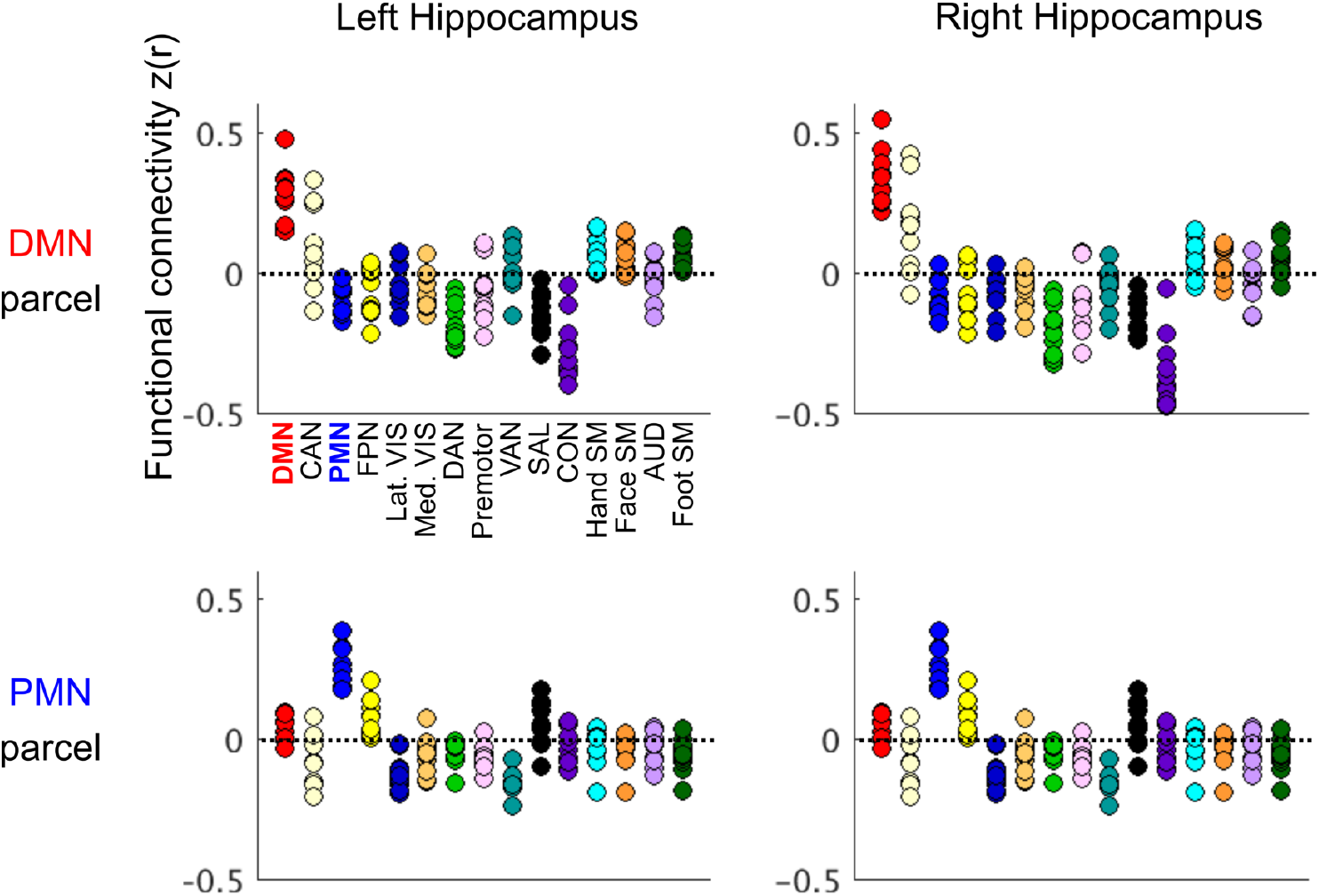
Mean functional correlation of WTA parcels to cortical resting state networks. The DMN (top row) and PMN (bottom row) parcels’ mean functional connectivity to all cortical resting state networks for the left and right hippocampus for each MSC subject. In addition to the parcels’ FC to its winning network, the mean FC to the runner-up networks (CAN & FPN for DMN & PMN respectively) can also be observed.

**Table S1.** Amount of variance explained in hippocampal RSFC by Gradient and Parcel factors. In an ANCOVA testing both the gradient (AP axis) and parcel factors head-to-head, the table depicts the amount of variance explained in FC difference (DMN - PMN) by one factor while accounting for the other for all MSC subjects and MSC06-Rep. The amount of variance explained by the factors alone are shown in **Figure 4** for the MSC subjects and in **Figure S3D** for MSC06-Rep.

| Subject | Gradient | Parcel |
| --- | --- | --- |
| MSC01 | 0.063 | 0.046 |
| MSC02 | **0.183** | 0.018 |
| MSC03 | **0.108** | 0.065 |
| MSC04 | 0.003 | **0.279** |
| MSC05 | 0.034 | **0.119** |
| MSC06 | **0.118** | 0.076 |
| MSC07 | **0.095** | 0.054 |
| MSC08 | 0.003 | **0.142** |
| MSC09 | 0.047 | **0.056** |
| MSC10 | **0.075** | 0.011 |
| MSC06-Rep | 0.061 | **0.090** |

**Figure S8.**
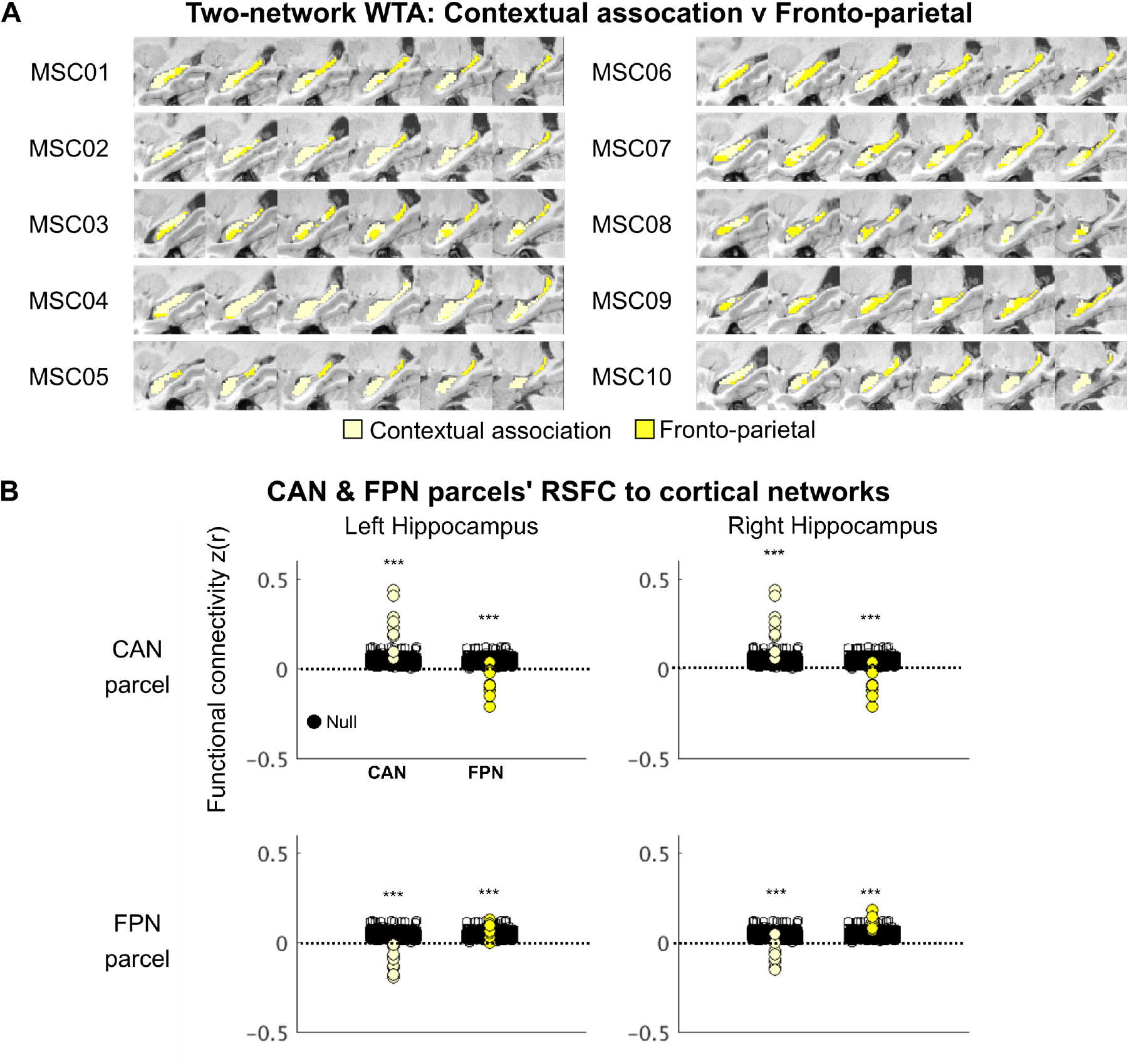
Significance testing of two-network (CAN & FPN) WTA parcellation of the hippocampus. We first defined the **(A)** WTA parcellation of the hippocampus using the CAN and FPN, which demonstrates an anterior-posterior axis of organization. The anterior hippocampus is connected to the CAN and the posterior to the FPN. Using the defined CAN and FPN parcels, we found that **(B)** the parcels’ RSFC to their winner networks is significantly different from a participant-specific null distribution.

**Figure S9.**
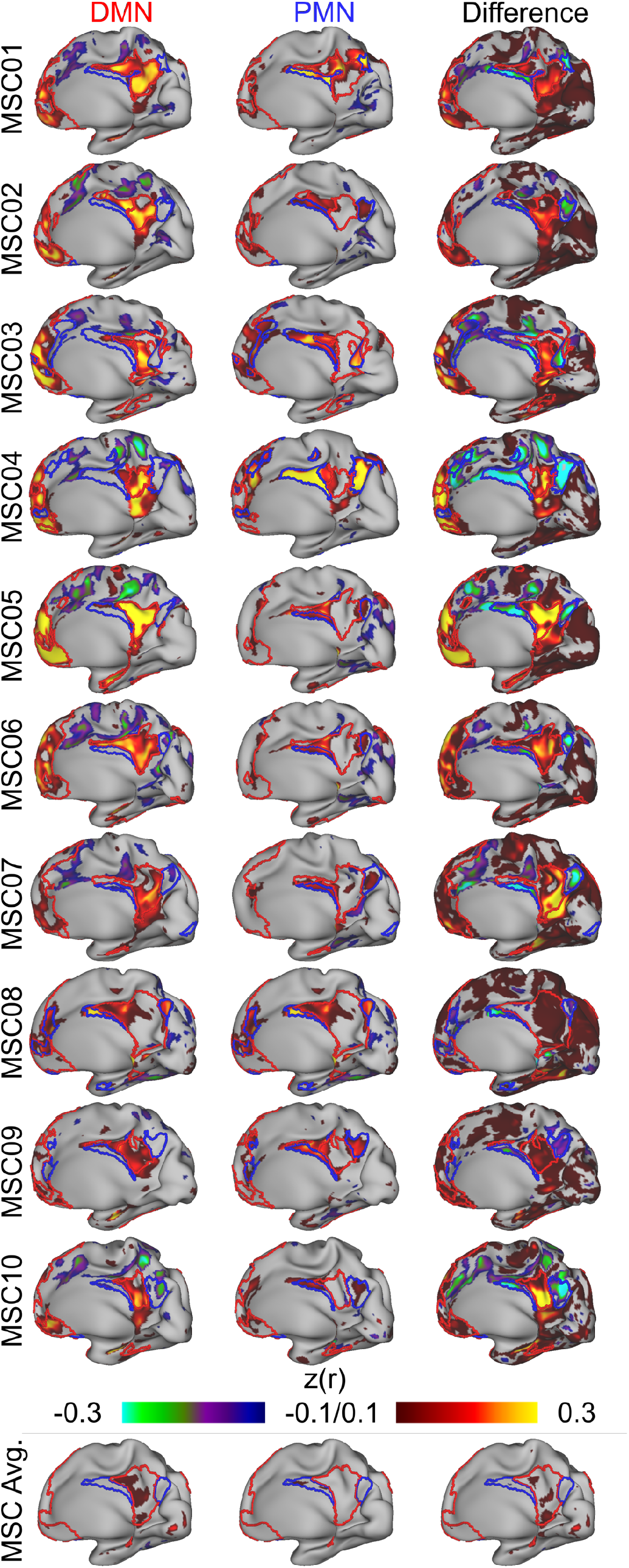
. Functional Connectivity of Individual-Specific Hippocampal DMN and PMN parcels for left hippocampus. Right hemisphere is shown. The first two columns show the connectivity patterns of the anterior, default mode network (DMN) and the posterior, parietal memory network (PMN) parcels in the hippocampus. The third column depicts the difference maps of functional connections for the right hippocampus. The color scale for the last column is represented such that the warm colors represent greater DMN correlation and the cool colors represent greater PMN correlation. **Figure 3** shows the difference maps of the right hippocampal functional connectivity to the cortex.

**Figure S10.**
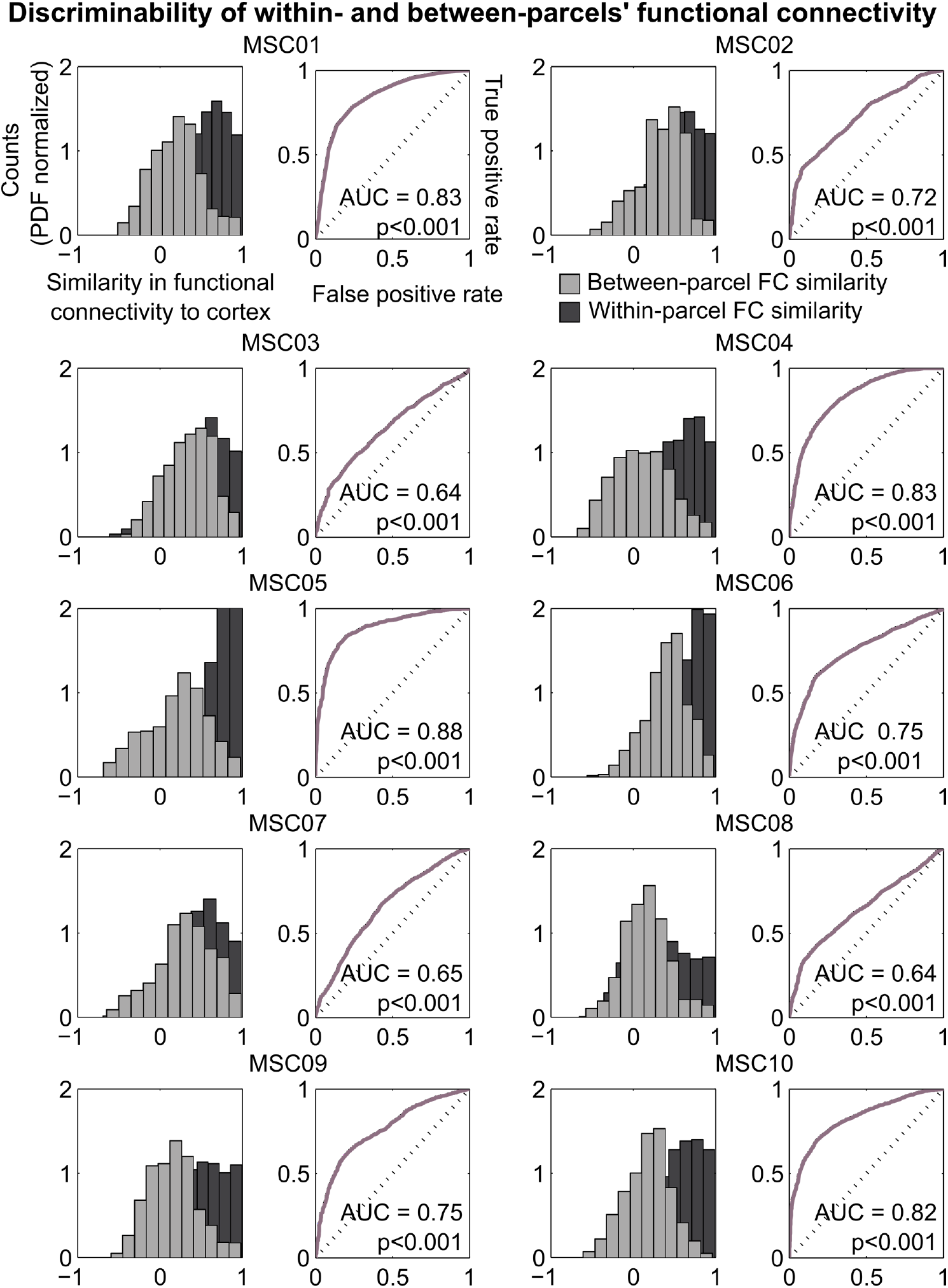
Discriminability of Within- and Between-parcels’ Functional Connectivity to the Cortex. The distribution of similarity in functional connectivity seed maps between pairs of voxels that are within the same hippocampal parcel vs. between different hippocampal parcels are discriminable as defined by a receiver-operator characteristic (ROC) curve. The area under the curve (AUC) represents the probability that an ideal observer would be able to adjudicate between these two distributions.

**Figure S11.**
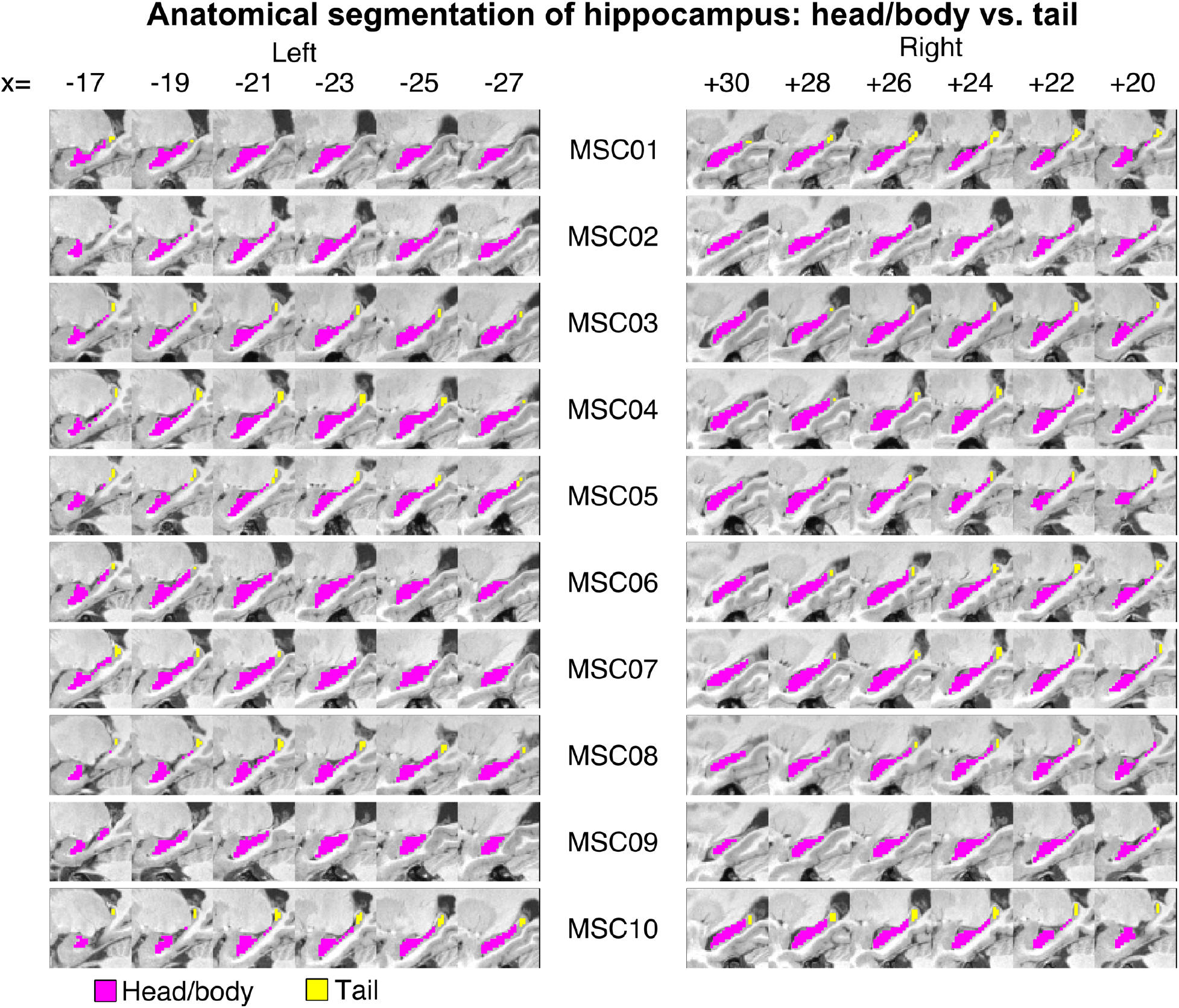
Anatomical segmentation of the hippocampus into head/body and tail. We used previously defined anatomical landmarks to segment the hippocampal body from the tail for each individual according to their unique hippocampal anatomy. Anatomical segmentations were then used to generate spatial correlation maps with the cortex in **Figure 5** as well as calculate spatial overlap between resting-state-derived functional parcellations and anatomical parcellations in **Figure S12**.

**Figure S12.**
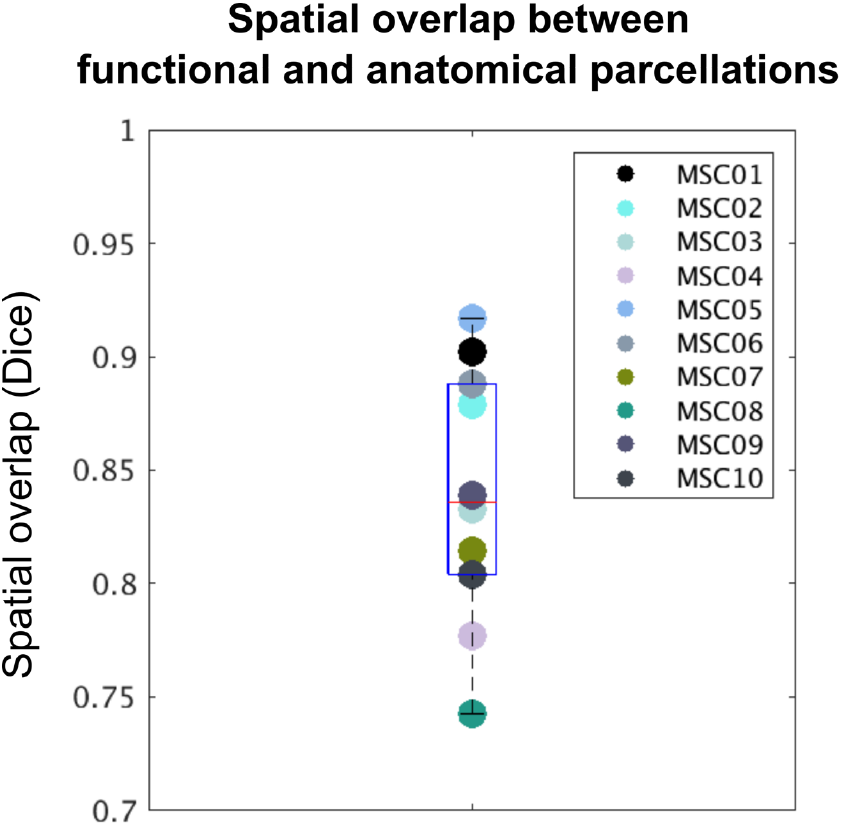
Spatial overlap between functionally-defined and anatomically-defined segmentations of the hippocampus. MSC subjects exhibit a high degree of spatial overlap (>0.7) between their subject-specific anatomical and WTA-defined parcels.

## Notes

https://openneuro.org/datasets/ds000224

### Competing Interest Statement

The authors have declared no competing interest.

